# Disruption of the MYC Super-Enhancer Complex by Dual Targeting of FLT3 and LSD1 in Acute Myeloid Leukemia

**DOI:** 10.1101/2022.01.17.476469

**Authors:** William M. Yashar, Brittany M. Curtiss, Daniel J. Coleman, Jake Van-Campen, Garth Kong, Jommel Macaraeg, Joseph Estabrook, Emek Demir, Nicola Long, Dan Bottomly, Shannon K. McWeeney, Jeffrey W. Tyner, Brian J. Druker, Julia E. Maxson, Theodore P. Braun

## Abstract

Mutations in Fms-like tyrosine kinase 3 (FLT3) are common drivers in acute myeloid leukemia (AML) yet FLT3 inhibitors only provide modest clinical benefit. Prior work has shown that inhibitors of lysine-specific demethylase 1 (LSD1) enhance kinase inhibitor activity in AML. Here we show that combined LSD1 and FLT3 inhibition induces synergistic cell death in FLT3-mutant AML. Multi-omic profiling revealed that the drug combination disrupts STAT5, LSD1, and GFI1 binding at the MYC blood super-enhancer, suppressing super-enhancer activation as well as MYC expression and activity. The drug combination simultaneously results in the accumulation of repressive H3K9me1 methylation, an LSD1 substrate, at MYC target genes. We validated these findings in 72 primary AML samples with the nearly every sample demonstrating synergistic responses to the drug combination. Collectively, these studies provide preclinical rationale for the investigation of dual FLT3/LSD1 inhibition in a clinical trial.

## INTRODUCTION

Mutations in Fms-like tyrosine kinase 3 (FLT3) occur in nearly a third of all patients with acute myeloid leukemia (AML) and are associated with an inferior overall survival (1). The most frequent mutation in FLT3 is the internal tandem duplication (ITD) of the juxta-membrane domain (2). While small molecule inhibitors of FLT3 kinase produce higher overall response rates and improved survival compared to salvage chemotherapy in patients with relapsed or refractory FLT3-ITD positive AML, FLT3 inhibitor monotherapy is rarely curative and responses are short-lived (3–5). There is a clinical need for approaches to deepen the initial response to FLT3 inhibitors, enabling longer-lasting clinical responses.

An approach to improving responses to FLT3 inhibitors in AML is to simultaneously target aberrant FLT3 activity and its downstream mediators. A major driver of mutant-FLT3-dependent oncogenesis is the MYC proto-oncoprotein (6,7). MYC, a critical regulator of proliferation and differentiation, is overexpressed in the vast majority of patients with AML (8). Reactivation of MYC-controlled oncogenic networks by the bone marrow microenvironment promotes FLT3 inhibitor resistance (8,9). These findings suggest that improved responses to FLT3 inhibitors may be achieved with combination strategies that target MYC-dependent proliferative programs.

Direct inhibition of MYC has been an objective of anti-cancer therapeutic development for over the last twenty years. However, MYC has been considered undruggable due to its intrinsically disordered nature and lack of enzymatic activity (10). Another approach is to instead disrupt the molecular mechanisms that drive *MYC* overexpression. In blood cells, *MYC* expression is regulated by a blood-specific super-enhancer complex (BENC), which is bound by numerous transcription factors and global chromatin activators (11,12). Recent studies in AML cell lines demonstrated that small molecule inhibitors targeting these activating chromatin complexes, including BRD4, resulted in a loss of *MYC* expression and leukemia cell death (13–16). However, initial clinical trials have only shown modest clinical activity and substantial toxicity (17).

An alternate approach is to simultaneously target two factors that regulate *MYC* gene expression. The chromatin regulatory protein lysine specific demethylase 1 (LSD1) is a well-established regulator of *MYC* gene expression. LSD1 regulates gene expression by the removal of activating methylation marks on lysine 4 of histone 3 (H3K4) and repressive methylation marks on lysine 9 of histone 3 (H3K9) or by the recruitment of repressive complexes to gene promoters (18,19). Inhibitors of LSD1 have been shown to decrease MYC in AML cell lines and primary samples (20–22). Our prior work and that of other groups shows that LSD1 inhibition augments the efficacy of kinase inhibitors in AML (21–23). However, the extent to which synergy exists between LSD1 and FLT3 inhibition and the underlying mechanism of drug synergy has not been investigated.

Here we report *ex vivo* drug screening data on a cell line model of FLT3-ITD positive AML and primary FLT3-ITD positive AML samples demonstrating that LSD1 inhibition potentiates the efficacy of FLT3 inhibition. Using high-sensitivity epigenetic profiling, we establish that FLT3/LSD1 inhibitors disrupt regulatory factor binding at the *MYC* BENC, resulting in a loss of *MYC* expression. Using short-term *ex vivo* culture, we confirm that these transcriptional and epigenetic responses to combined FLT3/LSD1 inhibition occur in primary FLT3-ITD positive leukemic blasts. Collectively, these data provide preclinical support for the clinical investigation of combined FLT3/LSD1 inhibition in patients with FLT3-ITD positive AML.

## RESULTS

### Combined FLT3/LSD1 inhibition synergistically represses MYC transcriptional programs, while activating PU.1 programs

Prior work from our lab and others suggest that the combination of kinase and LSD1 inhibition may be an effective therapeutic strategy in AML (20–23). To establish whether this approach is effective for FLT3-ITD AML, we treated FLT3-ITD-positive (MOLM13 and MV4-11) and FLT3-ITD-negative (K562) cell lines with multiple FLT3/LSD1 inhibitors. We observed potent synergy between the FLT3 inhibitors and LSD1 inhibitors in the FLT3-ITD-positive cell lines but not in the FLT3-ITD-negative cell lines, demonstrating that this drug combination has specificity for FLT3-ITD-positive AML (Fig. 1A-B; Supplementary Fig. 1). In addition, we observed that the drug combination increased both early (Annexin V+/PI-) and late (Annexin V+/PI+) apoptosis populations with minimal toxicity to healthy CD34+ cells (Supplementary Fig. 2A-C). These results indicate that synergy exists between FLT3/LSD1 inhibition in FLT3-ITD-positive AML.

**Fig. 1.**
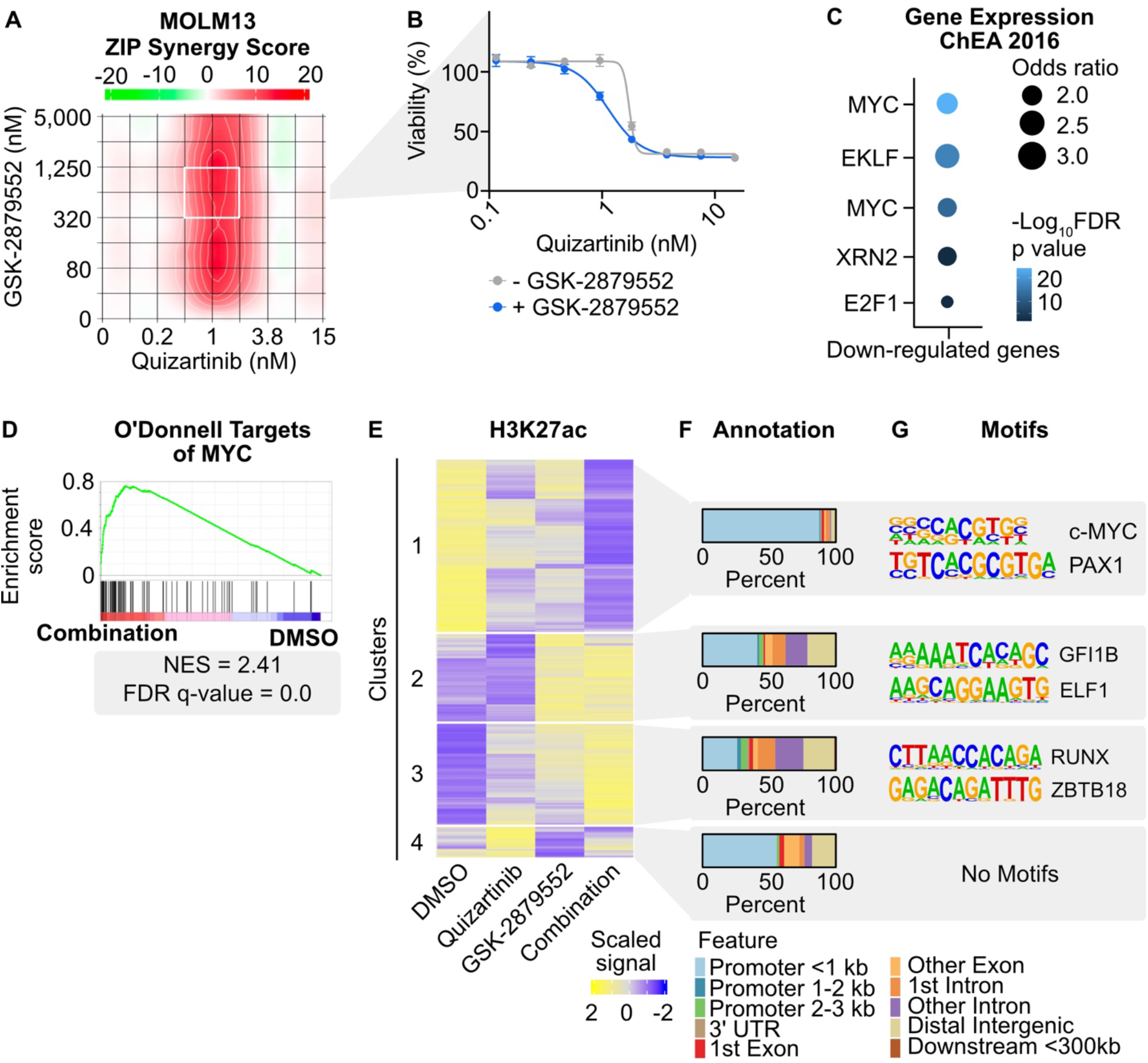
Transcriptional and chromatin dynamics in response to combined FLT3/LSD1 inhibition in FLT3-ITD-positive AML. **A**, MOLM13 cells were treated in triplicate with an 8×8 dose matrix of quizartinib and GSK-2879552 for 3 days prior to viability assessment by CellTiter Aqueous colorimetric assay. Zero interaction potency (ZIP) synergy scores were calculated on the average values for each drug dose. The white box indicates the quizartinib and GSK-2879552 concentrations corresponding to maximal synergy. **B**, Quizartinib response curves with and without GSK-2879552 (638 nM, which is the concentration corresponding to maximal synergy in a). The GSK-2879552 response curve with and without quizartinib is shown in Supplementary Fig. 1. **C**, MOLM13 cells were treated with quizartinib (1 nM), GSK-2879552 (100 nM), the combination, or an equal volume of DMSO vehicle for 24 hours prior to RNA isolation and RNA sequencing. GO analysis was performed on genes with synergistically decreased expression with the drug combination relative to DMSO. **D**, GSEA was performed comparing the drug combination to DMSO. **E**, MOLM13 cells were treated with quizartinib (1 nM), GSK-2879552 (100 nM), the combination, or an equal volume of DMSO vehicle for 2 hours prior to CUT&Tag for H3K27ac. Unsupervised hierarchical clustering of regions of differential read pileup with drug treatment. **F**, Annotation of peaks in clusters from (**D**). **G**, Motif enrichment for regions of differential H3K27ac read density. Top two *de novo* motifs with p <10^−12^ shown.

To understand the mechanism of synergy, we performed RNA-seq on MOLM13 cells treated with quizartinib, GSK-2879552, or the combination for 24 hours. Unsupervised clustering of differentially expressed genes revealed clusters of genes that were either synergistically upregulated or downregulated by the drug combination (Supplementary Fig. 2D; Supplementary Table 1). Transcription factor target gene enrichment analysis of the synergistically downregulated genes revealed an enrichment of MYC target genes (Fig. 1C; Supplementary Fig. 2E; Supplementary Table 2). In contrast, the genes synergistically upregulated by the drug combination were enriched for SPI1/PU.1 targets. The *SPI1* gene encodes the transcription factor PU.1, which is critical for coordinating myeloid differentiation (24). To further corroborate these gene expression profiles, gene set enrichment analysis (GSEA) was performed, which revealed depletion of MYC target genes in cells treated with the drug combination (Fig. 1D; Supplementary Table 3). Collectively, we observed that the combination activates an anti-proliferative and pro-differentiative transcriptional program with repression of MYC target genes and activation of PU.1 target genes.

### Combined FLT3/LSD1 inhibition disrupts chromatin dynamics at distinct genomic loci

A key component of LSD1 inhibitor activity has been ascribed to disruption of the LSD1 scaffolding function, loss of GFI1 binding to chromatin, and reactivation of enhancers associated with differentiation (25). Therefore, to characterize the early chromatin dynamics following combined FLT3/LSD1 inhibition, we utilized cleavage under targets and tagmentation (CUT&Tag) to assess changes in acetylation of histone 3 lysine 27 (H3K27ac), a marker of transcriptional activation, in MOLM13 cells 2 hours following drug treatment (26). Unsupervised clustering of the regions with differential H3K27ac signal revealed four clusters (Fig. 1E; Supplementary Table 4). The peaks in cluster 1 showed synergistic repression of acetylation by the drug combination and were localized primarily to promoters (Fig. 1F). The cluster 1 peaks were enriched for MYC motifs, consistent with the findings of decreased gene expression of MYC target genes (Fig. 1G). An example of a synergistically downregulated peak is observed at the PVT1 promoter, a known regulator of *MYC* expression (Supplementary Fig. 2F) (27). Cluster 2 contained peaks with increased signal largely driven by LSD1 inhibition. Peaks in this cluster were nearly equally distributed at promoter and non-promoter elements and were enriched for GFI1/GFI1B motifs. Cluster 3 peaks were also localized at promoter and non-promoter elements and showed enrichment for RUNX1 motifs. RUNX1 is a critical regulator of myeloid differentiation and potentiates the transcriptional activation activity of PU.1 (28). An example of synergistically upregulated enhancers from clusters 2 and 3 were observed upstream of the lysozyme promoter, which is expressed in mature granulocytes (Supplementary Fig. 2G). These data collectively show that the drug combination alters the chromatin landscape at distinct genomic loci. Furthermore, based on pathway analysis and motif enrichment, MYC, GFI1, RUNX, and PU.1 transcription factors are candidate regulators of these chromatin dynamics.

### Chromatin segmentation reveals that MYC-, STAT5-, and PU.1-driven molecular programs underlie the response to combined FLT3 and LSD1 inhibition

Our CUT&Tag results revealed substantial changes in histone acetylation at both promoters and outside promoters in response to combined FLT3/LSD1 inhibition, arguing that both types of regulatory elements have distinct roles in the drug response. We therefore profiled a series of covalent histone marks in MOLM13 cells 6 hours following drug treatment, enabling the segmentation of chromatin into promoters and enhancers. Trimethylation of histone 3 lysine 4 (H3K4me3) is primarily localized at promoters, whereas monomethylation of histone 3 lysine 4 (H3K4me1) is predominantly at enhancers (29). We also profiled H3K27ac at this same time point to understand transcriptional activation at prompters and enhancers (26). Following LSD1 inhibition, we observed regions of increased H3K4me3 and H3K4me1 levels consistent with the known demethylase activity of LSD1 for H3K4me1/3 (Supplementary Fig. 3A-D) (25,30). Unsupervised clustering of regions of differential H3K27ac signal at promoters and enhancers revealed multiple patterns of regulation (Supplementary Table 5). Similar to the global 2-hour acetylation CUT&Tag data, we observed a large cluster of repressed H3K27ac signal at promoters (cluster P2) that were enriched for MYC motifs and for GO terms associated with cell cycle and proliferation as well as a cluster of increased H3K27ac signal (cluster P1) enriched for GFI1/GFI1B motifs (Fig. 2A and B; Supplementary Fig. 3E; Supplementary Table 6). At enhancers, we identified a cluster of suppressed H3K27ac signal (cluster E3) associated with STAT5 motifs along with a cluster of increased H3K27ac signal (cluster E4) enriched for SPI1/PU.1 motifs (Fig. 2C and D; Supplementary Fig. 3F). This data suggests that combined FLT3/LSD1 inhibition simultaneously activates GFI1/GFI1B-bound promoters and represses the activation of MYC-bound promoters. In parallel, the drug combination suppresses STAT5-bound enhancers and activates enhancers bound by PU.1.

**Fig. 2:**
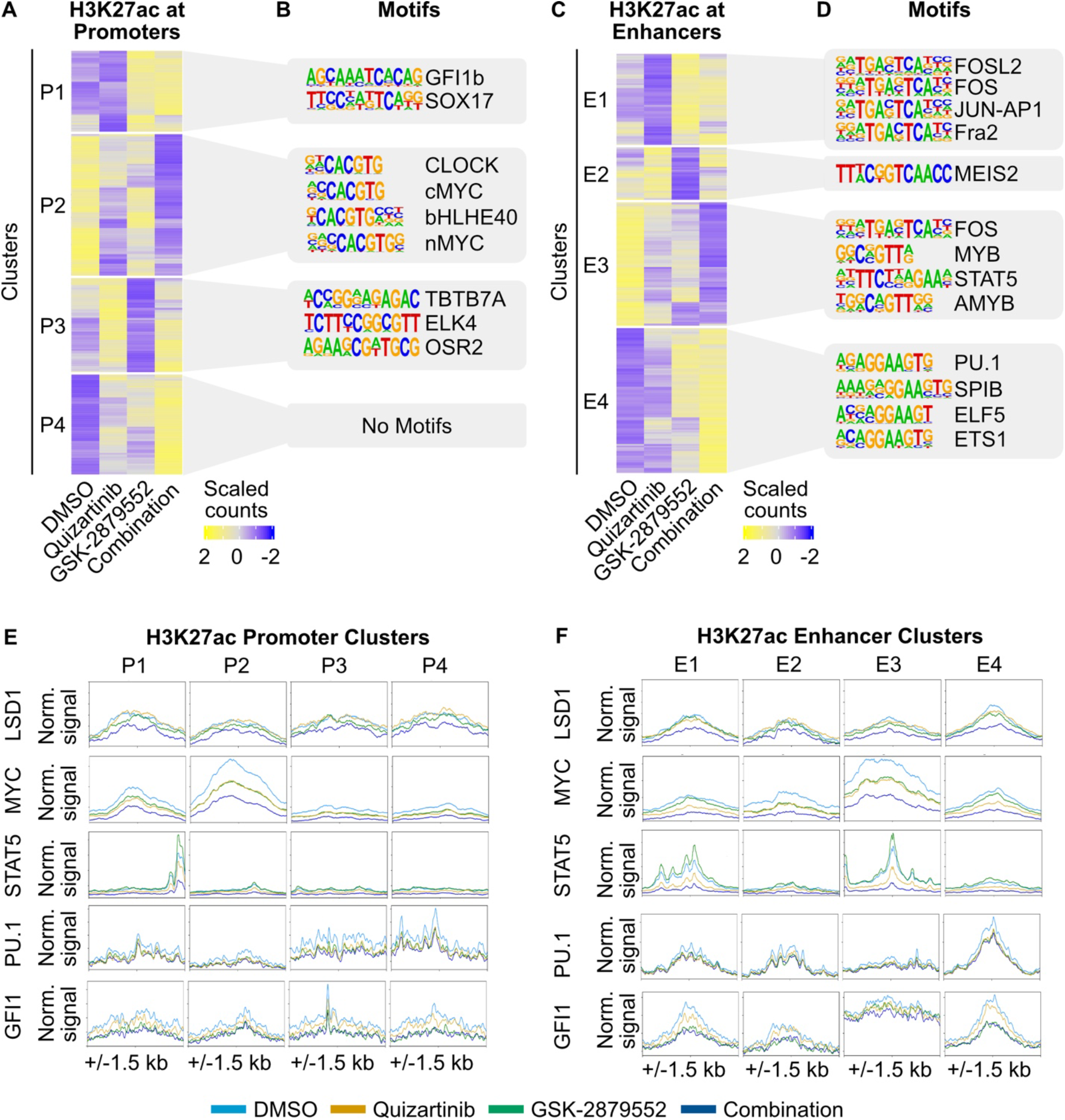
Discrete components of the response to FLT3/LSD1 inhibition are mediated by enhancers and promoters. **A**, MOLM13 cells were treated with quizartinib (1 nM), GSK-2879552 (100 nM), the combination, or an equal volume of DMSO vehicle for 6 hours prior to CUT&Tag for H3K27ac, H3K4me1, and H3K4me3. On the basis of these marks, chromatin was segmented into enhancers and promoters. Unsupervised hierarchical clustering of differential H3K27ac regions at promoters. **B**, Transcription factor motif enrichment is shown with top 4 known motifs. If no significant known motifs were identified, then *de novo* motifs with p < 10^−12^ are shown. **C, D**, Same analyses as (**A**) and (**B**) were performed at enhancers. **E**, MOLM13 cells were treated with quizartinib (1 nM), GSK-2879552 (100 nM), the combination, or an equal volume of DMSO for 6 hours. LSD1, MYC, and STAT5 binding was assessed by ChIP-seq. PU.1 and GFI1 binding was assessed by CUT&RUN. Transcription factor binding profiles at promoters with differential H3K27ac identified in (**A**). **F**, Transcription factor profiles at enhancers with differential H3K27ac identified in (**C**).

To better characterize the transcription factor regulators driving the response to combined FLT3/LSD1 inhibition, we profiled the genome-wide binding of multiple candidate transcription factors following single or dual drug treatment using chromatin immunoprecipitation sequencing (ChIP-seq), cleavage under targets and release using nuclease (CUT&RUN), and CUT&Tag. We then examined the pileup of these factors at each cluster of differential H3K27ac signal at promoters and enhancers (Fig. 2E and F; Supplementary Fig. 3G and H). We observed modest enrichment of LSD1 at all acetylated promoter and enhancer regions. MYC pileup was most pronounced at differential H3K27ac promoters (cluster P2), and enhancers (cluster E3) that were suppressed by the combination. In both clusters, a greater loss of MYC signal was observed with combined FLT3/LSD1 inhibition compared with no drug or single drug controls. STAT5 binding was localized to differential enhancers that were downregulated by quizartinib and/or enriched for STAT5 motifs (clusters E1 and E3), which is consistent with studies demonstrating that STAT5 is a primary downstream target of FLT3 inhibitors (31). Notably, combined FLT3/LSD1 inhibition resulted in a depletion of STAT5 signal at these clusters. PU.1 and GFI1 showed specific enrichment at differential enhancers with decreased H3K27ac signal following LSD1 inhibition (clusters E1 and E4). LSD1 inhibition led to a loss GFI1 signal at these clusters, consistent with previously reported displacement of GFI1 from chromatin upon LSD1 inhibition (25). While RUNX1 and CEBPA were enriched at both differential promoters and enhancers, they did not demonstrate appreciable changes in signal following drug treatment. Collectively these results implicate MYC, STAT5, PU.1, and GFI1 in the synergistic cytotoxicity of combined FLT3/LSD1 inhibition.

### Loss of *MYC* expression is critical for the response to combined FLT3/LSD1 inhibition

Our transcriptomic and epigenetic analyses nominated MYC as a key driver of the molecular responses to combined FLT3/LSD1 inhibition. We observed that combined FLT3/LSD1 inhibition results in the suppression in the total abundance of *MYC* transcript and in a genome-wide decrease in MYC binding (Fig. 3A and B). We confirmed that the transcriptional suppression of *MYC* was associated with a decrease in MYC protein abundance (Supplementary Fig. 4A). Modulation of MYC target gene expression can be influenced both by changes in *MYC* gene expression and activity. MYC regulates the transcription of cell cycle proteins through the recruitment of pause-released factors to poised RNA Polymerase II (RNA PolII) (32). We observed that the drug combination increased RNA PolII pausing at MYC-bound genes, indicating a disruption in the ability of MYC to promote RNA PolII pause release at its target genes (Fig. 3C; Supplementary Fig. 4B and C; Supplementary Table 7). In addition, we observed an increase in TP53 protein levels and an enrichment of a causal network controlled by TP53 following the drug combination, consistent with repression of MYC-dependent cell cycle regulation (Supplementary Fig. 4D; Supplementary Table 8). Together, these findings suggest a mechanism of combined FLT3/LSD1 that suppresses MYC expression and activity.

**Fig. 3:**
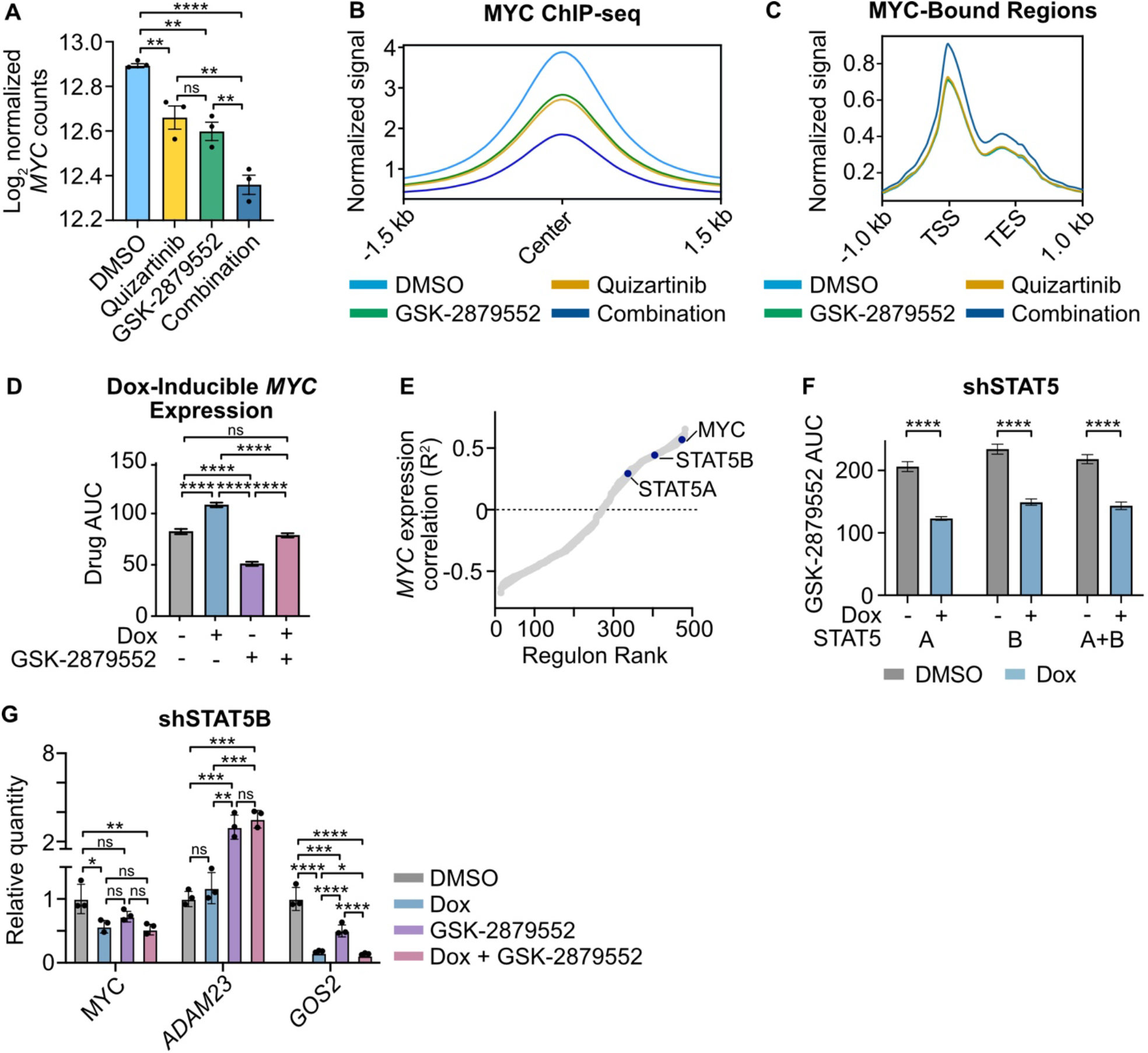
*MYC* expression is suppressed by combined FLT3/LSD1 inhibition and is associated with STAT5 regulatory activity. **A**, Normalized read counts for *MYC* from MOLM13 RNA-seq presented in Fig. 1. Statistical significance was determined by twoway ANOVA with a Holm-Śidák post-test correction. **B**, MYC binding profile at consensus peaks from MOLM13 ChIP-seq presented in Fig. 2. **C**, MOLM13 cells were treated with quizartinib (1 nM), GSK-2879552 (100 nM), the combination, or an equal volume of DMSO vehicle for 6 hours prior to CUT&Tag for RBP1. RBP1 binding profile at RBP1 and MYC co-bound regions. **D**, MOLM13 cells were transduced with lentiviral particles harboring a doxycycline-inducible *MYC* expression vector. Cells were treated with doxycycline (1 µg/mL) or DMSO for 48 hours and then plated in an 8×8 matrix of quizartinib and GSK-2879552 for 3 days prior to viability assessment by CellTiter Aqueous colorimetric assay. AUC data from the 311 nM GSK isoline (the concentration corresponding to maximal synergy in the *MYC* over-expressed MOLM13 cells in Supplementary Fig. 4) is shown. Dose responses and synergy over the entire drug matrix is shown in Supplementary Fig. 4. Statistical significance was determined by two-way ANOVA with a Holm-Śidák post-test correction. **E**, Spearman’s correlation of normalized *MYC* gene counts and regulon enrichment scores from baseline RNA-seq performed on patients in the BeatAML cohort. Regulons are ranked by goodness of fit (R^2^). **F**, MOLM13 cells were transduced with lentiviral particles harboring a doxycycline-inducible *STAT5* knockdown. GSK-2879552 AUC of cells treated with doxycycline (1 µg/mL) or DMSO for 72 hours. The GSK-2879552 response curves are shown in Supplementary Fig. 5. Statistical significance was determined by Student’s t-test. **G**, qPCR assessment of gene expression in MOLM13 cells expressing a doxycycline-inducible *STAT5B* shRNA. Cells were treated with doxycycline (1 µg/mL) for 72 hours prior to the addition of GSK-2879552 (100 nM) for 24 hours. Expression was normalized to *GusB* as an endogenous control. Statistical significance was determined by two-way ANOVA with a Holm-Śidák post-test correction. ns = not significant, * = p < 0.05, ** = p < 0.01, *** = p < 0.001, **** = p < 0.0001, TSS = transcription start site, TES = transcription end site.

To evaluate the importance of *MYC* expression to the mechanism of dual FLT3/LSD1 inhibition, we derived a MOLM13 cell line with a doxycycline-inducible *MYC* expression construct (Supplementary Fig. 4E). *MYC* overexpression not only resulted in decreased sensitivity to the drug combination, but also attenuated the induction of apoptosis by the drug combination (Fig. 3D; Supplementary Fig. 4F-I). These data suggest that forced expression of *MYC* partially abrogates the effect of combined FLT3/LSD1 inhibition.

To identify potential regulators of *MYC* gene expression in AML that may mediate the response to the combination, we performed a regulon enrichment analysis using RNA-seq on 681 primary AML samples. We generated a regulon enrichment scores for each sample that reflected the predicted activity for 468 different transcription factors. Correlation of these transcription factor activity scores with *MYC* gene expression revealed a strong positive correlation with STAT5 regulon activity (Fig. 3E; Supplementary Table 9). As FLT3 is a known activator of STAT5, we generated MOLM13 cell lines with doxycycline-inducible knockdown of *STAT5* to evaluate its role in the response to combined FLT3/LSD1 inhibition (Supplementary Fig. 5A-C) (31). Knockdown of *STAT5A* and/or *STAT5B* resulted in increased sensitivity to GSK-2879552 (Fig. 3F; Supplementary Fig. 5D-F). Furthermore, we observed synergy between *STAT5* knockdown and GSK-2879552, demonstrating that a loss of STAT5 activity is sufficient to recapitulate a portion of the quizartinib effect (Supplementary Fig. 5G-J). *STAT5* knockdown resulted in reduced expression of *MYC* as well as the MYC target genes *ADAM23* and *GOS2*, which was potentiated by the presence of GSK-2879552 (Fig. 3G; Supplementary Fig. 5K and L) (33,34). Collectively, these results suggest that the STAT5-MYC axis plays a major role in the response to dual FLT3/LSD1 inhibition.

### Quizartinib suppresses STAT5 binding to the MYC blood super-enhancer

To identify the mechanism by which STAT5 regulates *MYC* expression in FLT3-ITD AML, we examined the binding data from our STAT5 ChIP-seq data. While we did not identify a STAT5 binding event at the MYC promoter, ranking STAT5 peaks by normalized read pileup revealed strong binding event at the MYC BENC consistent with previous findings (Fig. 4A; Supplementary Tables 10, 11) (11,35). Notably, the MYC BENC was among the differentially downregulated H3K27ac enhancers where STAT5 ChIP-seq signal was depleted following combined FLT3/LSD1 inhibition (cluster E3). STAT5-bound elements within the MYC BENC showed a rapid decrease in H3K27ac signal after treatment with quizartinib or the drug combination (Fig. 4B). To characterize the changes in activity of the MYC BENC we performed assay for transposase-accessible chromatin with highthroughput sequencing (ATAC-seq) (27). This analysis revealed a loss of chromatin accessibility across all modules in response to drug combination treatment (Fig. 4C-E). Evaluation of STAT5 binding and H3K27ac signal revealed several sub-modules that also display dynamic behavior in response to drug treatment but have not been previously characterized (A0.1-0.5, C1, G1). Collectively, our findings nominate the MYC BENC as a crucial locus for downregulation of *MYC* gene expression by dual FLT3/LSD1 inhibition.

**Fig. 4:**
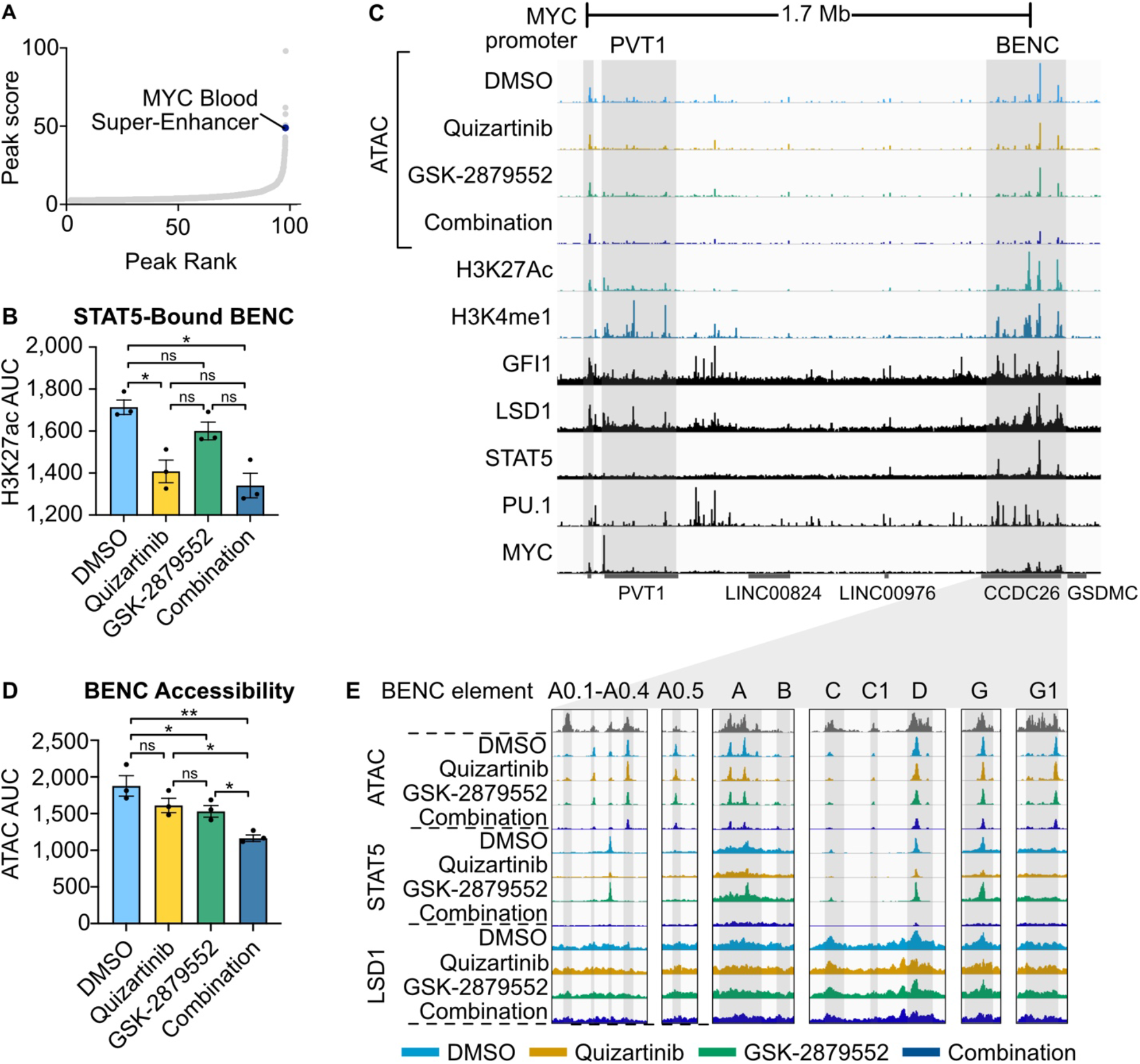
FLT3-Inhibition represses *MYC* expression through a loss of STAT5 binding to the MYC blood super-enhancer cluster. **A**, STAT5 peak read pile up ranked by overall signal strength in MOLM13 cells from Fig. 2. **B**, H3K27ac signal AUC at STAT5-bound BENC elements. H3K27ac signal data is from MOLM13 cells in Fig. 1 whereas the STAT5 signal data is from MOLM13 cells in Fig. 2. Statistical significance was determined by two-way ANOVA with a Holm-Šidák post-test correction. **C**, ATAC-seq was performed on MOLM13 cells quizartinib (1 nM), GSK-2879552 (100 nM), the combination, or an equal volume of DMSO vehicle for 24 hours. Representative histone modification and transcription factor tracks (from DMSO-treated MOLM13 cells in Fig. 2) shown at the extended *MYC* locus. **D**, ATAC signal AUC at all MYC BENC modules. Statistical significance was determined by two-way ANOVA with a Holm-Šidák post-test correction. **E**, ATAC, STAT5 ChIP-seq, and LSD1 ChIP-seq signal at twelve independent BENC modules. ns = not significant, * = p < 0.05, ** = p < 0.01, *** = p < 0.001, **** = p < 0.0001.

### LSD1 inhibition represses the expression of MYC and its target genes by altering GFI1/CoREST dynamics and histone modification

Our data demonstrates that dual FLT3/LSD1 inhibition suppresses *MYC* gene expression by displacement of STAT5 binding from the MYC BENC. However, the MYC BENC is bound by many other transcription factors, indicating that drug combination efficacy may be dependent on interruption of other MYC BENC-bound factors. Prior studies have shown that LSD1 inhibitor monotherapy decreases *MYC* expression (20–22). A critical component of LSD1-inhibitor efficacy is the disruption of LSD1 scaffolding of GFI1 from the CoREST transcription repressor complex (25). Examination of our GFI1 CUT&RUN data confirmed that GFI1 is not only bound to the MYC BENC, but also revealed a decrease in GFI1 signal at MYC BENC modules C and G following drug combination treatment (Fig. 5A). To evaluate the importance of GFI1 in the response to combined FLT3/LSD1 inhibition, we generated MOLM13 cell lines with doxycycline-inducible knockdown of *GFI1* (Supplementary Fig. 6A). We found that *GFI1* knockdown increased sensitivity to FLT3 inhibition and enhanced FLT3-inhibitor-dependent repression of *MYC* and its target genes (Fig. 5B and C; Supplementary Fig. 6B and C). Collectively, this data indicates that displacement of GFI1 binding by LSD1 inhibition reduces *MYC* expression and is important to combined FLT3/LSD1 inhibitor response.

**Fig. 5:**
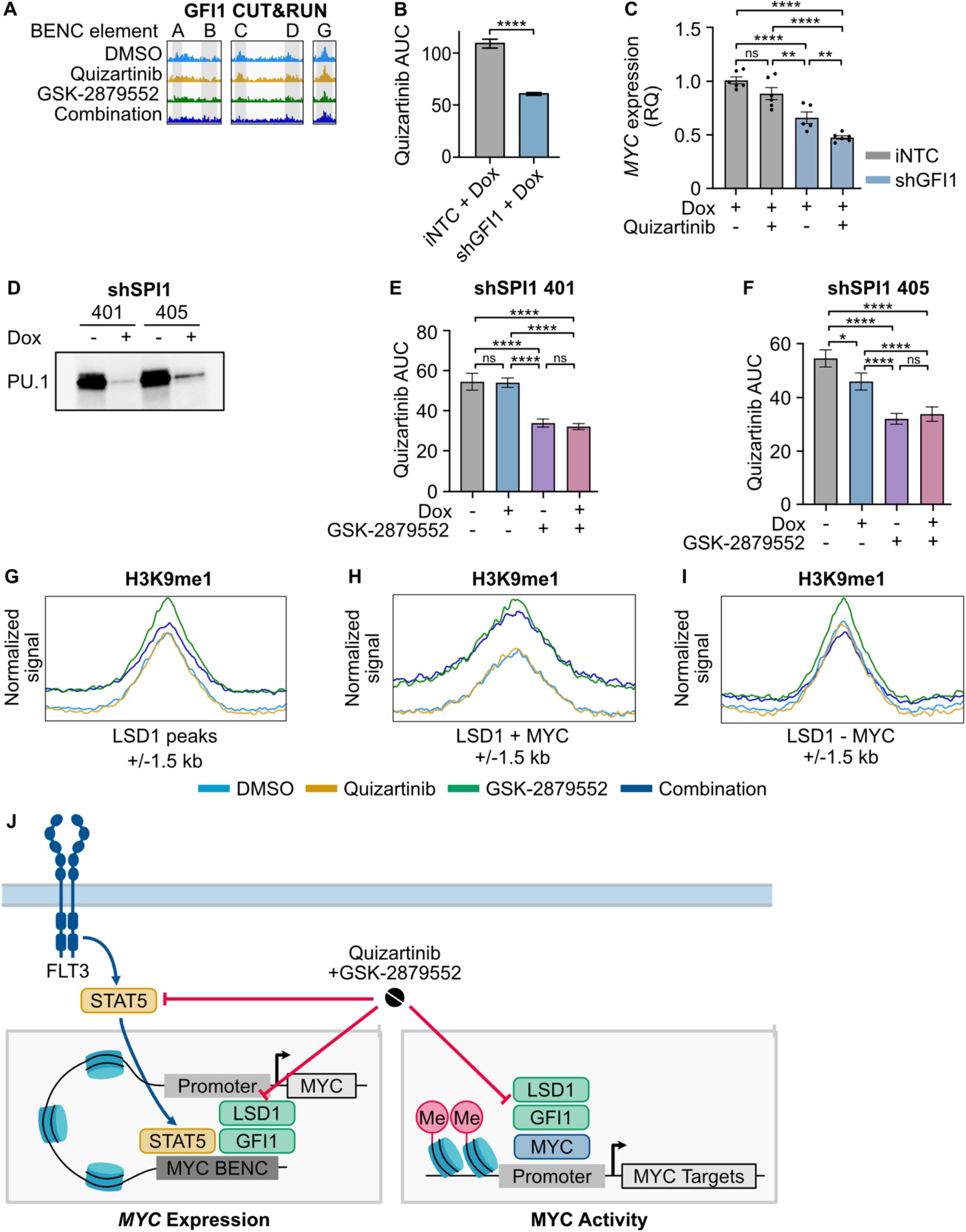
LSD1 inhibition disrupts GFI1/CoREST binding at the MYC BENC and induces a gain of H3K9me1 binding at MYC-bound promoters. **A**, GFI1 CUT&RUN signal from Fig. 2 at five BENC modules. **B**, MOLM13 cells were transduced with lentiviral particles harboring a doxycycline-inducible non-targeting codon (iNTC) or *GFI1* knockdown vector. Quizartinib AUC of cells treated with doxycycline (1 µg/mL) or DMSO for 72 hours. Substantial knockdown was observed in the absence of doxycycline treatment, so only doxycycline-treated samples were compared. The quizartinib response curves are shown in Supplementary Fig. 6. Statistical significance was determined by Student’s t-test. **C**, qPCR assessment of gene expression in cells treated with doxycycline (1 µg/mL) for 72 hours prior to the addition of quizartinib (1 nM) for 24 hours. Expression was normalized to *GusB* as an endogenous control. Statistical significance was determined by two-way ANOVA with a Holm-Śidák post-test correction. **D**, MOLM13 cells were transduced with lentiviral particles harboring a doxycycline-inducible *SPI1* knockdown vector. Western blot for PU.1 in cells treated with doxycycline (1 µg/ml) or an equivalent volume of DMSO for 72 hours. **E, F**, Cells were treated with doxycycline (1 µg/mL) or DMSO for 48 hours and then plated in an 8×8 matrix of quizartinib and GSK-2879552 for 3 days prior to viability assessment. AUC data from the 311 nM GSK isoline (the concentration corresponding to maximal synergy in the *SPI1* knockdown MOLM13 cells) is shown. Dose responses and synergy over the entire drug matrix is shown in Supplementary Fig. 6. Statistical significance was determined by two-way ANOVA with a Holm-Śidák post-test correction. **G-I**, MOLM13 cells were treated with quizartinib (1 nM), GSK-2879552 (100 nM), or the combination for 6 hours prior to CUT&Tag for H3K9me1. Normalized signal for H3K9me1 at LSD1-bound regions, LSD1 and MYC co-bound regions, and at regions bound by LSD1 but not MYC. **J**, Model describing the drug combination mechanism. Ns = not significant, * = p < 0.05, ** = p < 0.01, *** = p < 0.001, **** = p < 0.0001.

Our data demonstrate that dual FLT3/LSD1 inhibition exert a portion of their activity via a STAT5- and GFI1-dependent decrease in *MYC* gene expression. However, *MYC* over-expression only partially reverses the impact of combination treatment, suggesting the involvement of additional mechanisms. Prior work has shown that LSD1 inhibitor efficacy is also dependent on activation of PU.1-bound enhancers (36). Our transcriptional and epigenetic data shows that combined FLT3/LSD1 inhibition results in the activation of PU.1 target genes and acetylated enhancers enriched for PU.1 motifs. Therefore, we assessed the impact of *SPI1* (the gene that encodes the PU.1 protein) knockdown on the efficacy of combined FLT3/LSD1 inhibition. PU.1-deficient cells demonstrated no significant reduction in drug synergy, suggesting that PU.1 is not crucial for the efficacy of dual FLT3 and LSD1 inhibition (Fig. 5D-F; Supplementary Fig. 6D-I). In other cell types, LSD1 plays a role in gene activation via removal of repressive H3K9me1/2 marks (37). To evaluate this possible mechanism, we profiled the genome-wide distribution of H3K9me1 via CUT&Tag (Fig. 5G-I; Supplementary Fig. 6J). Inhibition of LSD1, with or without inhibition of FLT3, resulted in an accumulation of H3K9me1 at MYC target genes co-bound with LSD1. This was accompanied by a loss of the reciprocal activating mark acetylated histone 3 lysine 9 (H3K9ac), consistent with the observed decrease in the expression of MYC target genes. These findings suggest that dual inhibition of FLT3/LSD1 exerts locus-specific effects on the chromatin landscape by interrupting STAT5 and GFI1/CoREST transcriptional regulation as well as altering the balance of repressive H3K9 marks at MYC-bound promoters (Fig. 5J).

### Efficacy of combined FLT3/LSD1 inhibition in primary AML samples

We assessed the efficacy of combined FLT3/LSD1 inhibition on 72 primary AML samples using a 3-day *ex vivo* drug assay. Nearly every sample (94%; 68 of 72 samples) demonstrated a synergistic increase in efficacy of dual agent therapy over single agents alone (Fig. 6A and B; Supplementary Fig. 7A and B; Supplementary Table 12). Although synergy was observed regardless of FLT3 mutation status, the mean quizartinib AUC was lower in FLT3-ITD-positive samples (140.1) as compared to FLT3-wildtype samples (172.4) or samples harboring a FLT3 mutation other than FLT3-ITD (168.2). To characterize the determinants of response to combined FLT3/LSD1 inhibition, we performed a regulon enrichment analysis on baseline RNA sequencing performed on the cohort (Supplementary Table 13). MYC and STAT5B regulon activity were among the strongest correlates with the degree of combination synergy (Fig. 6C). CDK4, a known transcriptional target of MYC, was the strongest correlate (38). These findings reveal that AML samples with high baseline MYC activity have the greatest sensitivity to combined FLT3/LSD1 inhibition.

**Fig. 6:**
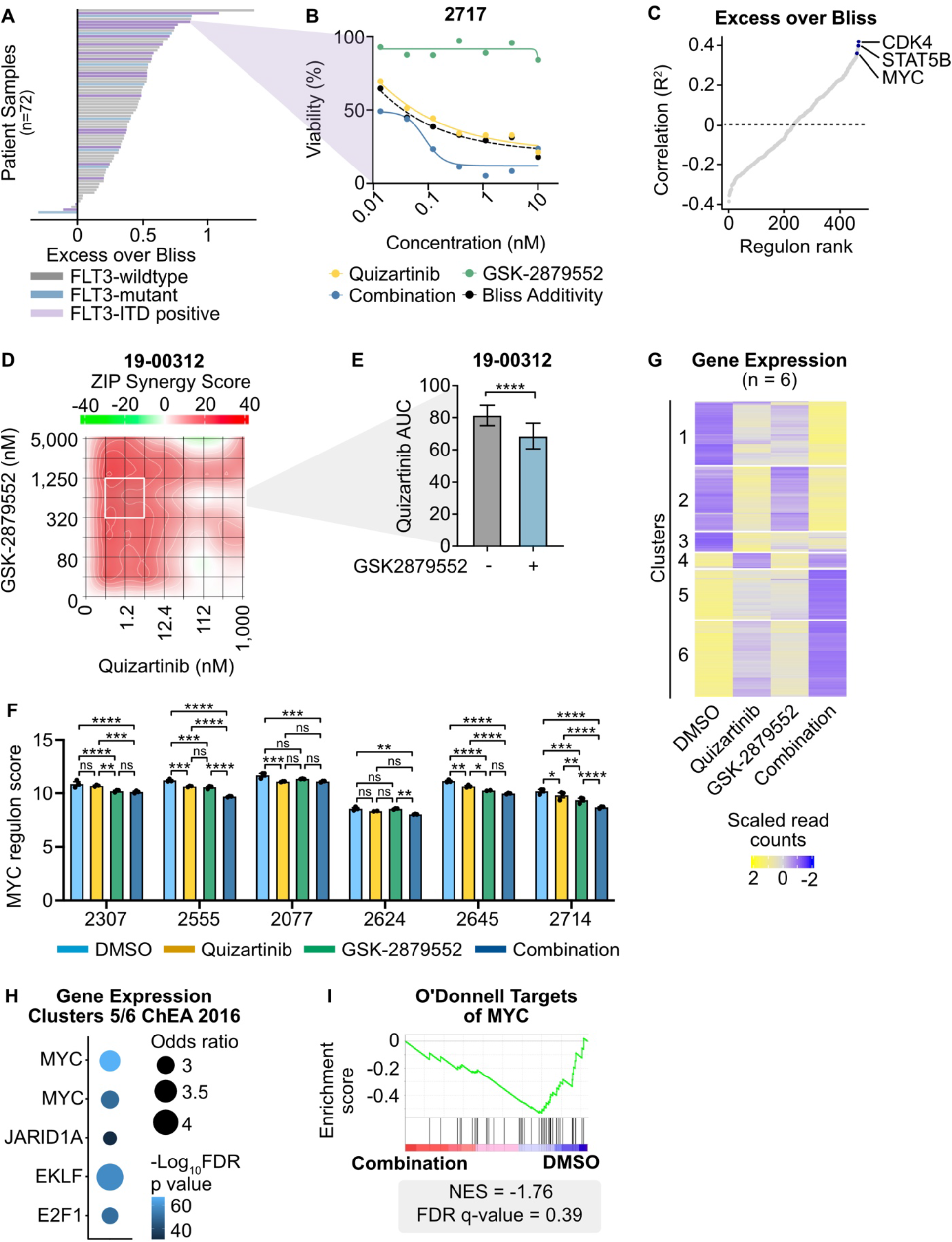
Combined FLT3/LSD1 inhibition drives synergistic cell death by repressing a MYC-dependent transcriptional network in primary AML blasts. **A**, Primary AML blasts from 72 total samples (18 FLT3-ITD-positive) were cultured for 3 days along a 7-point curve with either quizartinib, GSK-2879552, or equimolar amounts of the drug combination. Cell viability was assessed by CellTiter Aqueous colorimetric assay. The excess over bliss score was calculated using cell viability at corresponding drug concentrations. Each bar represents the mean excess over bliss across all concentrations. Bar color indicates FLT3 mutation status. **B**, Dose response curves for quizartinib, GSK-2879552, and the drug combination in a FLT3-ITD-positive AML sample from (**A**). **C**, Spearman’s correlation of excess over bliss and regulon enrichment scores. **D**, Primary blasts from a FLT3-ITD-positive AML sample were treated in triplicate with an 8×8 dose matrix of quizartinib and GSK-2879552 for 3 days prior to viability assessment by CellTiter Aqueous colorimetric assay. ZIP synergy scores were calculated on the average values for each drug dose. **E**, AUC data from the 628 nM GSK isoline (the concentration corresponding to maximal synergy in d) is shown. Statistical significance was determined by Student’s t-test. **F**, Bulk RNA-seq was performed on six independent, FLT3-ITD-positive patient samples treated in triplicate with 500 nM quizartinib, 500 nM GSK-2879552, both drugs in combination, or an equivalent volume of DMSO for 24 hours. Normalized MYC regulon enrichment scores were calculated from RNA-seq. Statistical significance was determined by two-way ANOVA with a Holm-Śidák post-test correction. **G**, Unsupervised hierarchical clustering of genes differentially expressed upon drug treatment. **H**, Transcription factor target enrichment from clusters in (**G**). **I**, Select GSEA plot from clusters in (**G**). ns = not significant, * = p < 0.05, ** = p < 0.01, *** = p < 0.001, **** = p < 0.0001.

To characterize the response to combined FLT3/LSD1 inhibition in patient samples, we performed drug sensitivity studies and RNA-seq on 6 FLT3-ITD-positive patient samples treated with single or dual agent therapy. Similar to the pattern of synergy observed in cell lines, we found drug synergy across a broad range of doses and synergistic induction of apoptosis in primary AML blasts (Fig. 6D and E; Supplementary Fig. 7C and D). MYC regulatory activity was downregulated all samples by the drug combination, although differing patterns of individual drug effect were observed (Fig. 6F; Supplementary Fig. 7E). Unsupervised hierarchical clustering of differentially expressed genes across all 6 samples revealed a similar pattern to that observed in MOLM13 cells (Fig. 6G; Supplementary Table 14). Transcription factor target analysis revealed suppression of MYC target genes and activation of SPI1/PU.1 target genes (Fig. 6G; Supplementary Fig. 7F; Supplementary Table 15). Finally, GSEA revealed the drug combination decreased expression of MYC target genes and increased expression of differentiation-associated genes (Fig. 6I; Supplementary Fig. 7G; Supplementary Table 16). Collectively, this data confirms the findings of MYC gene expression and regulon activity from AML cell lines in primary AML blasts.

### Dual FLT3/LSD1 inhibition suppresses MYC super-enhancer activity in primary AML blasts

Our initial investigations in AML cell lines demonstrate that suppression of chromatin accessibility at the MYC super-enhancer is an important mechanism of cytotoxicity produced by combined FLT3/LSD1 inhibition. To validate this mechanism in primary AML blasts, we performed single cell ATAC-seq on primary AML blasts from three FLT3-ITD-positive patients following 24 hours of treatment with combined quizartinib and GSK-2879552 or DMSO. Clustering and cell type identification by CD34+ expression revealed that the samples were largely comprised of myeloid blasts (Fig. 7A-C; Supplementary Fig. 8A-C). Separation of cell clusters based on treatment revealed population shifts in response to treatment (Fig. 7D-F; Supplementary Fig. 8D-F). Assessment of read pileup at the MYC BENC showed that these population shifts were associated with decreased chromatin accessibility at most BENC modules (Fig. 7G-J; Supplementary Table 17). However, each sample showed a distinct pattern of change. Sample 2684 showed decreases in the majority of modules, 2645 demonstrated the most prominent decreases in modules A and G, and sample 2603 exhibited the strongest decreases in modules F and G. Ranking of peaks in sample 2684 by log_10_ fold change in peak score between the drug combination and DMSO revealed that module C was amongst the regions with the greatest decrease in accessibility (Fig. 7K). Collectively, these results demonstrate that suppression of MYC BENC activity is a conserved feature of the response to dual FLT3/LSD1 inhibition in primary AML samples. However, patient-to-patient heterogeneity does exist, suggesting diversity in the regulatory factors that sustain MYC BENC activity in primary AML samples.

**Fig. 7:**
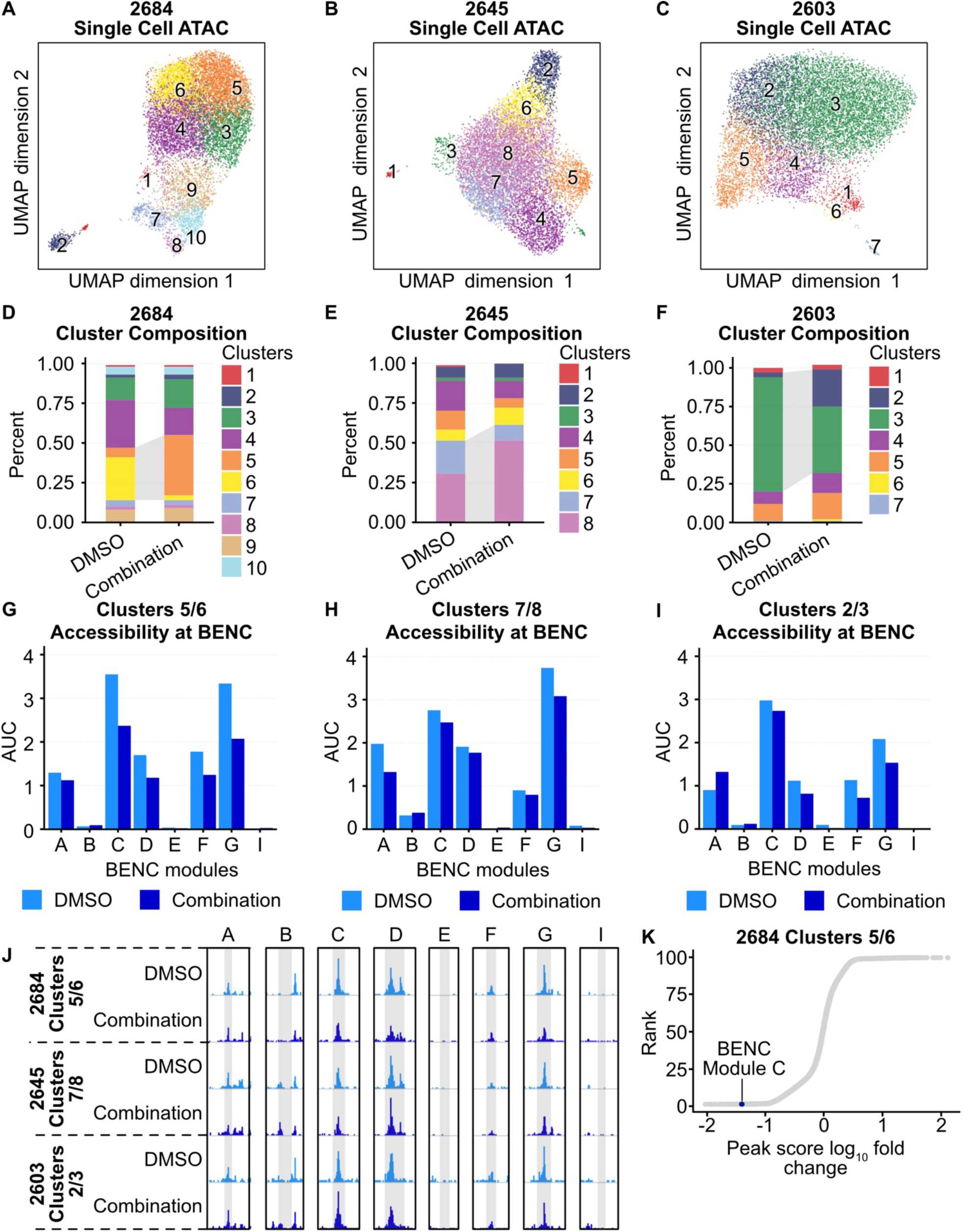
Dual FLT3/LSD1 inhibition results in a shift from a MYC super-enhancer-high to a MYC super-enhancer-low cell state in primary AML blasts. **A-C**, Single cell ATAC-seq was performed on three AML patient samples with DMSO or quizartinib (500 nM) and GSK-2879552 (500 nM) for 24 hours. UMAP of DMSO and drug-treated cells colored by cluster. **D-F**, Proportions of DMSO-treated and drug-treated cells assigned to each cluster. Dynamic clusters were identified as the populations that shift between DMSO and drug-treated conditions and with greater accessibility at CD34. Dynamic clusters are highlighted with gray shading between bars. **G-I**, AUC of accessibility at each BENC module is displayed. **J**, Treatment group pseudo-bulked accessibility at the MYC BENC modules separated by treatment and patient sample. **K**, Ranking by log_10_(peak score fold change) between DMSO-treated and the combination-treated cells within dynamic clusters.

## DISCUSSION

Activating mutations in FLT3 are amongst the most common molecular events in AML (1). While FLT3 inhibitors are clinically available, they produce only modest improvements in survival (4,5). Here, we demonstrated that LSD1 inhibition potentiates the efficacy of FLT3 inhibition in FLT3-ITD AML cell lines and primary cell blasts. High-resolution transcriptomic and epigenetic profiling revealed that the mechanism of synergy is in part due to depletion of regulatory transcription factor binding, STAT5 and GFI1, at the MYC BENC. Moreover, we identified additional evidence that dual FLT3/LSD1 inhibition results in the accumulation repressive H3K9me1 marks at MYC-controlled proliferation genes. These findings provide rationale for the clinical investigation of combined FLT3/LSD1 inhibitors for patients with FLT3-ITD AML and reveals how epigenetic therapies augment the activity of kinase inhibitors in AML.

A crucial component to the mechanism of FLT3/LSD1 inhibitor synergy was altering *MYC* expression through regulation of the MYC BENC. While others have demonstrated that *MYC* transcription can by altered by inhibiting regulators of this super-enhancer, regulation of MYC BENC activity by combined epigenetic modulatory drugs and kinase inhibition is a novel approach to targeting this central oncogenic regulator (13–16). Our single cell ATAC-seq analysis revealed substantial variation in the pattern of MYC BENC module utilization between AML samples at baseline and in response to drug treatment (Fig. 7). Indeed, other studies have suggested that each BENC module is bound by a distinct set of transcription factors and regulates *MYC* expression in specific blood cell lineages (11). How MYC BENC module use varies between molecularly defined AML subtypes and how this relates to drug responses is an important question for future studies.

Prior work on LSD1 inhibitors has largely implicated the pro-differentiation effects of these drugs as the central mechanism of cytotoxicity. Our work here shows that LSD1 inhibition strongly activates enhancers that are bound by PU.1. Other groups have shown that suppression of *SPI1* expression results in a block in LSD1-inhibitor-induced differentiation and decreased cytotoxicity (36). While our work confirmed the role of PU.1 as a putative mediator of LSD1-inhibitor responses, we found that *SPI1* knockdown had little effect on the transcriptional or cytotoxic response to dual FLT3/LSD1 inhibition (Fig. 5; Supplementary Fig. 6). It is unclear whether the pro-differentiation transcriptional effects observed in our study are important to the drug effect. Investigation of the pro-differentiation effects of dual FLT3/LSD1 inhibition will be an important question for future investigation.

Previous studies of LSD1 inhibitors have also demonstrated that drug efficacy is dependent on the interruption of LSD1 scaffolding activity rather than its demethylation activity (20,39). Our work confirmed that a critical component of LSD1 inhibitor activity is the disruption of LSD1 binding to GFI1/CoREST (Fig. 5; Supplementary Fig. 6). However, LSD1 inhibition also resulted in the accumulation of repressive H3K9me1 marks at the promoters of MYC target genes. While LSD1 canonically demethylates activating H3K4 marks, alternative LSD1 complexes remove repressive H3K9 methylation marks in cells from other tissues, resulting in transcriptional activation (37,40). In prostate cancer, LSD1 forms a chromatin-associated complex with androgen receptor that demethylates H3K9 and de-represses androgen receptor target genes. In neuronal cells, on the other hand, an LSD1 isoform, LSD1+8a, complexes with supervillain (SVIL) and demethylates H3K9me2 to regulate neuronal differentiation. Further work needs to be done to nominate binding factors with LSD1 or LSD1 isoforms that, as a complex, functions as a transcriptional activator by H3K9 demethylation.

Collectively, our work demonstrates that LSD1 inhibition enhances the activity of FLT3 inhibition in FLT3-ITD AML. The efficacy of the drug combination is dependent on the simultaneous disruption of STAT5 and GFI1 from the MYC blood super-enhancer complex, resulting in repressed *MYC* expression, as well as the accumulation of repressive H3K9me1 at MYC target genes.

## MATERIALS AND METHODS

### Cell and Patient Sample Culture

#### Cell Lines

MOLM13 cells (DSMZ) were cultured in RPMI (Gibco) supplemented with 20% fetal bovine serum (FBS, HyClone), 2 mM GlutaMAX (Gibco), 100 units/mL Penicillin, and 100 µg/mL Streptomycin (Gibco). MV-4-11 and K562 cells (ATCC) were cultured in IMDM supplemented with 10% fetal bovine serum (FBS, HyClone), 2 mM GlutaMAX (Gibco), 100 units/mL Penicillin, and 100 µg/mL Streptomycin (Gibco). All cells were cultured at 5% CO_2_ and 37°C. Cell lines were tested monthly for mycoplasma contamination.

#### Patient Samples

All patients gave informed consent to participate in this study, which had the approval and guidance of the institutional review boards at Oregon Health & Science University (OHSU), University of Utah, University of Texas Medical Center (UT Southwestern), Stanford University, University of Miami, University of Colorado, University of Florida, National Institutes of Health (NIH), Fox Chase Cancer Center and University of Kansas (KUMC). Samples were sent to the coordinating center (OHSU; IRB#9570; #4422; NCT01728402) where they were coded and processed. Mononuclear cells were isolated by Ficoll gradient centrifugation from freshly obtained bone marrow aspirates or peripheral blood draws. Clinical, prognostic, genetic, cytogenetic, and pathologic lab values as well as treatment and outcome data were manually curated from patient electronic medical records. Genetic characterization of the leukemia samples included results of a clinical deep-sequencing panel of genes commonly mutated in hematologic malignancies (Sequenome and GeneTrails (OHSU); Foundation Medicine (UTSW); Genoptix; and Illumina). Patient samples were cultured in RPMI with 10% FBS and 10% HS-5 conditioned media (ATCC) or SFEMII supplemented with 1x StemSpan CD34+ Expansion Media and 1 µM UM729 (StemCell Technologies).

#### Colony Assay

Whole bone marrow was obtained (AllCells) and CD34+ cells were selected using CD34 MicroBead Kit (Miltenyi Biotec) according to manufacturer’s instructions. For the colony assay, 500 CD34+ cells were used per replicate and plated in MethoCult™ H4435 Enriched (StemCell Technologies). The four groups were treated with quizartinib (1 nM), GSK-2879552 (100 nM), the combination, or DMSO. Plates were incubated for 14 days in 5% CO_2_ and 37°C. Samples were imaged using STEMvision (StemCell Technologies) and blinded prior to counting by another investigator by assigning letters randomly. ImageJ (NIH) was used to count colonies after blinding.

#### Drug Synergy

Drug synergy was assessed using an 8 × 8 matrix of drug concentrations. Cells were treated for 72 hours prior to MTS assay to evaluate viability. Cell viability was used to calculate drug synergy with SynergyFinder based on the ZIP reference model (41).

#### RNA Interference

Two SMARTvector Inducible shRNAs for Human SPI1 (V3IHSHER_10431275, V3IHSHER_10642739), two for STAT5A (V3IHSHEG_6691183, V3IHSHEG_4988581) two for STAT5B (V3IHSHER_4778243, V3IHSHER_6411380) and two for GFI1 (V3IHSHER_5266412 and V3IHSHER_5697821) in a hEF1a-TurboRFP or hEF1a-TurboGFP (STAT5A) backbone were obtained from Horizon Discovery. Both SPI1 shRNA construct showed effective knockdown of SPI1. STAT5A V3IHSHEG_6691183 was specific for STAT5A, STAT5A V3IHSHEG_4988581 was ineffective against either STAT5A or B, STAT5B V3IHSHER_6411380 was selective for STAT5B and V3IHSHER_4778243 knocked down both STAT5A and B. GFI1 V3IHSHER_5266412 produced effective GFI1 KD while V3IHSHER_5697821 was ineffective. Lentivirus was produced by transfecting Lenti-X 293T cells (Clontech) with the SMARTvector transfer plasmid and packaging/pseudotyping plasmids. psPAX2 was a gift from Didier Trono (Addgene plasmid # 12260; http://n2t.net/addgene:12260; RRID:Addgene_12260) and pMD2.G was a gift from Didier Trono (Addgene plasmid #12259; http://n2t.net/addgene:12259; RRID:Addgene_12259). The supernatants containing lentivirus was collected after 48 hours of culture and filtered with a 0.45 um filter. MOLM13 cells were transduced with virus via spinnoculation in the presence of polybrene. Transduced cells were selected with 1 µg/mL puromycin to produce a stable cell line.

#### MYC *Overexpression*

For human *MYC* overexpression, pDONR223_MYC_WT was a gift from Jesse Boehm & Matthew Meyerson & David Root (Addgene plasmid # 82927; http://n2t.net/addgene:82927; RRID:Addgene_82927) and cloned into pCW57.1, a gift from David Root (Addgene plasmid #41393; http://n2t.net/addgene:41393; RRID:Addgene_41393). Lentiviral particles were generated as above and MOLM13 cells were selected after viral transduction with 1 µg/mL puromycin. After selection, cells were treated with 1 µg/mL doxycycline for 72 hours prior to experiments.

#### Apoptosis

Apoptosis was assessed 48-72 hours after drug treatment by flow cytometry using an eBioscience Annexin V-APC apoptosis detection kit (ThermoFisher) according to the manufacturer’s instructions.

### Sequencing Methods

#### Bulk RNA-Seq

MOLM13 cells were treated with 1 nM quizartinib, 100 nM GSK-2879552, the combination, or equal volume of DMSO for 24h. Total RNA was isolated using a RNeasy Plus Mini Kit (Qiagen). BGI performed the library preparation and sequencing with 50BP SE sequencing. Patient samples were cultured in 10% HS-5 CM/RPMI with 20% FBS and treated with 500 nM quizartinib, 500 nM GSK-2879552, the combination, or the equivalent volume of DMSO for 24 hours. Total RNA was isolated with RNeasy Micro kit (Qiagen) according to the manufacturer’s instructions. Libraries were prepared using the NEBNext Low Input RNA Library Prep Kit for Illumina (NEB) according to the manufacturer’s instructions. Libraries were sequenced by the OHSU Massively Parallel Sequencing Shared Resource (MPSSR) using 100 BP SE sequencing on an Ilumina NovaSeq S1 flow cell.

#### Bulk ATAC-Seq

MOLM13 cells were treated with 1 nM quizartinib, 100 nM GSK-2879552, the combination, or an equivalent volume of DMSO for 24 hours. After treatment, 50,000 cells per replicate were harvested for Fast-ATAC sequencing performed as previously described (42). In brief, cells were resuspended in cold PBS and tagmentation master mix (25 ul of 2x tagmentation buffer, 2.5 ul of TDE1 [Illumina], 0.5 ul of 1% digitonin; 2x tagmentation buffer: 66 mM Tris-Acetate, pH 7.8, 132 mM potassium acetate, 20 mM magnesium acetate, 32% v/v N,N-Dimethylformamide) was added. Samples were incubated at 37°C for 30 minutes. DNA was purified using Zymo Clean and Concentrator 5 Kit (Zymo). Transposed DNA was amplified and purified as described previously with adapted primers (43,44). Samples were quantified using Qubit dsDNA HS Assay Kit (Invitrogen), pooled, and sequenced by BGI using 50 BP PE sequencing.

#### CUT&Tag

MOLM13 cells were treated with 1 nM quizartinib, 100 nM GSK-2979552, the combination, or an equal volume of DMSO for 2 or 6 hours. Benchtop CUT&Tag was performed as previously described (45). In brief, cells were counted, harvested, and centrifuged for 5 min at 300xg at room temperature. Cells were washed 2X in 1.5 mL wash buffer (20 mM HEPES pH 7.5, 150 mM NaCl, 0.5 mM Spermidine, 1x Protease inhibitor cocktail). Concanavalin A magnetic coated beads (Bangs Laboratories) were activated in binding buffer by washing 2X (20 mM HEPES pH 7.5, 10 mM KCl, 1 mM CaCl_2_, 1 mM MnCl_2_). Washed cells were separated into 100,000 cell aliquots and 10 ul of activated beads were added to each sample. Samples were rotated end-over-end for 7 minutes at room temperature. A magnetic stand was used to separate beads and the supernatant was removed. Primary antibody was diluted 1:50 in antibody buffer (20 mM HEPES pH 7.5, 150mM NaCl, 0.5 mM Spermidine, 1× Protease inhibitor cocktail, 0.05% digitonin, 2 mM EDTA, 0.1% BSA). The primary antibodies used were: H3K27ac (ab4729, Abcam), H3K4me1 (#5326, Cell Signaling Technologies), H3K4me3 (#9751, Cell Signaling Technologies), RBP1 (#2629, Cell Signaling Technologies), H3K9me1 (#C15410065, diagenode), H3K9ac (#C15410004, diagenode), CEBPA (#8178, Cell Signaling Technologies), and Normal Rabbit IgG (#2729, Cell Signaling Technologies as previously described was diluted 1:100 in dig-300 buffer (20 mM HEPES pH 7.5, 300 mM NaCl, 0.5 mM Spermidine, 1× Protease inhibitor cocktail, 0.01% digitonin) and added to samples (45). Samples incubated for 1 hour at room temperature on nutator. Samples were washed 2X with dig-300 buffer then resuspended in tagmentation buffer (dig-300 buffer with 1 mM MgCl_2_). Samples were incubated at 37°C for 1 hour. DNA was extracted with phenol:chloroform extraction. Samples were amplified by PCR using custom Nextera primers at 400 nM and NEBNext (46). PCR conditions were set to: 72°C for 5 minutes, 98°C for 30 seconds, 14 cycles of 98°C for 10 sec, 63°C for 10 sec, and 72°C for 1 minute. Libraries were purified with AMPure Beads (Beckman) and sequenced by the OHSU MPSSR on an Ilumina NovaSeq using 50 BP SE sequencing or NextSeq 500 using 37 BP PE sequencing.

#### CUT&RUN

MOLM13 cells were treated with 1 nM quizartinib, 100 nM GSK-2979552, the combination, or an equal volume of DMSO 24 hours. CUT&RUN was performed as previously described (47). Briefly, concanavalin A magnetic coated beads (Bangs Laboratories) were washed 2x in binding buffer (20 mM HEPES pH 7.5, 10 mM KCl, 1 mM CaCl_2_, 1 mM MnCl_2_) to activate. 500,000 cells per replicate were washed 2x with wash buffer (20 mM HEPES pH 7.5, 150 mM NaCl, 0.5 mM Spermidine, 1x Protease inhibitor cocktail). Cells were bound to beads by nutating for 10 minutes at room temperature. Cells were permeabilized and incubated overnight at 4°C on nutator with primary antibody in antibody buffer (wash buffer, 0.001% digitonin, 3 mM EDTA). The following antibodies were used at 1:50 PU.1 (MA5-15064, Invitrogen), GFI1 (ab21061, Abcam), and Normal Rabbit IgG (#2729, CST). Bead slurry was washed 2x with dig wash buffer (wash buffer, 0.001% dig) and resuspended with dig wash buffer and 1x pAG-MNase (Epicypher). Cell were incubated for 10 minutes on nutator at room temperature then washed 2x with dig wash buffer followed by resuspension in pAG-MNase reaction mix (dig wash buffer, 2 mM CaCl_2_). Bead slurry was incubated for 2 hours at 4°C on nutator. STOP buffer (340 mM NaCl, 20 mM EDTA, 4 mM EGTA, 50 µg/mL RNase A, 50 µg/mL glycogen, 0.02% dig) was then added, then tubes were incubated at 37°C for 10 minutes. DNA was extracted using phenol:cholorform extraction. Libraries were prepared using NEBNext Ultra II DNA Library Prep Kit (NEB), modified for CUT&RUN as previously described (48). After adapter ligation fragments were cleaned up with 1.75x AMPure beads (Beckman). Following PCR amplification, libraries were purified 2x with 1.2x AMPure beads to rid of adaptor fragments. Libraries were quantified on the 2100 Bioanalyzer instrument (Agilent) with the High Sensitivity DNA Analysis Kit (Agilent). Libraries were pooled and sequenced by MPSSR on a NextSeq 500 sequencer (Illumina) using 37 BP PE sequencing.

#### ChIP-Seq

ChIP-seq was performed using the SimpleChIP plus Enzymatic Chromatin IP Kit (Cell Signaling Technology). For each replicate, 2 × 10^7^ cells were fixed in 4% formaldehyde (Sigma) for 10 minutes at room temperature then quenched with glycine, washed and stored at −80°C until use. Nuclei were extracted according to the manufactures instructions and treated with 1.25 uL MNase in 500 uL Buffer B at 37°C for 20 minutes. Samples were sonicated on a Qsonic sonicator at 50% amplitude for 5 cycles of 15 sec on 15 sec off on ice. Crosslinks were reversed on a small aliquot of extracted chromatin quantified by OD_260_. A total of 5 µg of chromatin was used for each immunoprecipitation. The following antibodies were used with a total of 5 µg of each primary antibody used per immunoprecipitation: Rabbit anti-LSD1 (ab17721, Abcam), Rabbit ani-MYC (13987, Cell Signaling Technology), Rabbit anti RUNX1 (ab23980), Rabbit anti-STAT5 (94205S, Cell Signaling Technology) and Rabbit IgG (2729, Cell Signaling Technology). After overnight incubation, complexes were captured using protein G beads. Crosslinks were reversed and libraries prepped using an NEBNex Ultra II for DNA Library Prep kit. Libraries were sequenced by the OHSU MPSSR. The STAT5 libraries were sequenced by Genewiz using a HiSeqX and 150 BP PE sequencing.

#### Reverse Phase Protein Array (RPPA)

MOLM13 cells were treated for 24 hours with 1 nM quizartinib, 100 nM GSK-2979552, the combination. Cells were washed 2x in PBS then flash frozen. Cell pellets were lysed and processed by the University of Texas MD Anderson Cancer Center Functional Proteomics RPPA Core Facility.

#### Single Cell ATAC-Seq

Patient samples were treated with 500 nM quizartinib and 500 nM GSK-2879552 or an equal volume of DMSO for 24 hours. Nuclei were prepared using the demonstrated protocol for primary cell nuclei extraction from 10x Genomics. ATAC libraries were prepared using Chromium Single Cell ATAC Library and Gel Bead kit v1.1 (10x Genomics, 1000176). Libraries were sequenced with 50 BP PE sequencing by the OHSU MPSSR.

### Primary AML Blast Dataset

Gene mutation, drug response, and gene expression data from primary AML blasts was accessed through the BeatAML database (1). Samples from collected from patients who were in remission or had MDS at the time of collection were excluded from downstream analysis. Samples with quizartinib/GSK-2979552 single and dual agent drug response data were selected and stratified by FLT3-ITD mutation status. Gene expression dataset was downloaded in the form of RPKM.

### Data Analysis

#### Bulk RNA-Seq

Raw reads were trimmed with Trimmomatic and aligned with STAR (49,50). Two MOLM13 replicates were identified as outliers during QC and excluded from downstream analysis. Differential expression analysis was performed using DESeq2 (51). Raw p values were adjusted for multiple comparisons using the Benjamini-Hochberg method. GO analysis was performed using Enrichr and Gene Set Enrichment Analysis (GSEA) (52,53).

#### Regulon Enrichment Analysis

Regulon enrichment signatures were derived by comparing all gene expression features to the median expression level of all samples. Weights were assigned to the regulatory network and the positive and negative edges of each regulator were rank ordered. The first component of the enrichment signature, the local delta concordance signature, was derived by calculating the concordance between the magnitude of the weight assigned to a particular edge and the ranked position of that edge. The features associated with activation (positive edges within the regulatory network) were monotonically ranked from most lowly to highly expressed, whereas the features associated with repression (negative edges within the regulatory network) were ranked by a monotonically decreasing function. The second component of the enrichment signature, the local enrichment signature, was derived by capturing positional shifts in the local gene ranked signature and integrating those shifts with the weights assigned to overlapping features for a given regulon and the expression data set. The last component of the enrichment signature was derived by projecting the rank-sorted local signature onto the globally ranked feature space. A global enrichment signature from this projection was calculated for each regulator. The median of robust quantile-transformed ranked positions was used as the enrichment scores for both the local enrichment and global enrichment signatures. The median of robust quantile-transformed ranked positions for the three individual signatures were integrated and reported as single enrichment score for each regulator.

#### Bulk ATAC-Seq Analysis

ATAC libraries were aligned to the human genome (hg38) using BWA-MEM and sorted using SAMtools (54,55). Duplicates were marked with Sambamba, duplicates and mitochondrial reads were removed (56). Counts per million (CPM) normalized tracks were generated with deepTools (57). Differential accessibility was assessed with DESeq2 (51). GO analysis was performed using Genomic Regions Enrichment of Annotations Tool (GREAT) (58). CPM normalized tracks were merged with bigWigMerge and visualized using Integrative Genomics Viewer (59).

#### CUT&Tag and CUT&RUN Analysis

CUT&Tag and CUT&RUN libraries were aligned to the human genome (hg38) using Bowtie2 and the following options --local --very-sensitive-local --no-unal --no-mixed --no-dis- cordant --phred33 -I 10 -X 700 (60). Peaks were called using GoPeaks and the following option -mdist 1000 (61). High confidence peaks were defined by presence in 2/2 or 2/3 replicates. Consensus bed files were formed by merging the high confidence peaks from DMSO, quizartinib, GSK-2979552, and the combination using BEDTools (62). Differential peaks were identified using DESeq2 with default parameters (51). Heatmaps were produced using the ComplexHeatmap package from Bioconductor (63). Peaks were annotated to the nearest feature using ChIPseeker (64). GO analysis was performed using GREAT (58). Counts tables for differential peaks were produced using multicov from BEDTools (62). CPM normalized tracks, global signal heatmaps, and plot profiles at specified regions were generated using deepTools (57). Active promoters were defined by the presence of H3K4me3 and H3K27ac within 1000 bp of a TSS. Active enhancers were defined by the presence of H3K4me1 and H3K27ac beyond 1000 bp of a TSS. CPM normalized tracks were merged with bigWigMerge and visualized using Integrative Genomics Viewer (59). Read pileup within 1500 kb upstream and downstream of LSD1- bound peaks as well as those that include or exclude MYC and PU.1 was quantified using computeMatrix reference-point from deepTools (57). Read pileup signal was scaled and visualized with a heatmap using the ComplexHeatmap package from Bioconductor (63). RNA PolII pause indices were calculating by generating counts tables for each gene in a window from −30 to +100 around each TSS and then from the remainder of the gene bodies. Counts were divided by the region length and the pause index was calculated by dividing the TSS values by those for the corresponding gene body. Spearman correlation of transcription factor binding sites was assessed using multiBigwigSummary and plotCorrelation from deepTools (57).

#### ChIP-Seq

ChIP-seq libraries were aligned to the human genome (hg38) using bowtie using the following options -v 2 -m 1 --best –strata. Peaks were called using MACS2 callpeak with the -narrow option and IgG library used as the control (65). Bigwigs for signal visualization were generated by sequentially converting alignments to bed files, big bed files then CPM normalized bigwig files using UCSC Kent Utilities. Read pileup at enhancers and promoters was evaluated using deepTools (57). The STAT5 super-enhancer-like analysis was performed using the HOMER findPeaks -super command (66).

#### Reverse Phase Protein Array (RPPA)

The University of Texas MD Anderson Cancer Center Functional Proteomics RPPA Core Facility linearized, normalized, and batch-corrected (Level 4) the phosphoproteomic data. The data was analyzed with CausalPath (v 1.2.0) using a significant change of mean value transformation, an FDR threshold of 0.2, and 100 permutations for significance (67). The causal network was visualized on the CausalPath web server causalpath.org.

#### Drug Synergy

Excess over bliss was calculated from a Bliss independence model using base R and visualized with ggplot2 (v 3.3.5) (68). Bliss additivity was inferred from GSK-2879552 and quizartinib cell viability values using a Bliss Independence conditional response curve (68). The R stats package (v 3.6.2) was used to calculate the spearman correlation between the degree of excess over bliss and regulon enrichment scores.

#### Single Cell ATAC-Seq

Single cell ATAC-seq libraries were aligned to the human genome (hg38) using 10x Genomics Cell Ranger (69). ArchR was used in R to generate Arrow files for each fragment file, using a TSS enrichment of four and 5,000 unique fragments as cutoffs (70). An ArchRProject was created for each patient sample. Latent semantic indexing (LSI) was preformed using the addIterativeLSI function and clusters were added using the addClusters function. The UMAP for each patient sample, with both drug conditions integrated, was generated. The proportion of cells from each treatment condition was calculated and the dynamic CD34+ clusters were identified. Pseudo-bulk replicates from the dynamic clusters were normalized to TSS enrichment and used to generate bigwigs. The bigwigs were used to produce deepTools (57) matrices to assess accessibility at the MYC BENC modules. The bar plots were generated using ggplot2. Peaks were called on pseudo-bulk replicates from the dynamic clusters using the ArchR (70) addReproduciblePeakSet command and MACS2 (65). Dynamic cluster peaksets from the same patient were combined using BEDTools intersect –loj command (62). Peaks were ranked by their peak score and the log_10_(peak score fold change) between dynamic clusters was calculated using a custom script.

### Quantification and Statistical Analysis

In all cases, unless otherwise states, values are represented as the mean and error bars are the SEM. Prism software (version 9.1; Prism Software Corp.) or RStudio was used to perform statistical analysis, which are described in the figure legends. All data were analyzed with Student’s t-test or a two-way ANOVA followed by Holm-Šidák correction. For differential analysis of RNA-seq, CUT&Tag, CUT&RUN, and ATAC-seq p values were adjusted for repeated testing with a false discovery rate using the Benjamini-Hochberg method.

### Data Availability

All raw and processed sequencing data generated in this study are available upon request.

## AUTHORS’ DISCLOSURES

**W.M. Yashar** is a former employee of Abreos Biosciences, Inc. and was compensated in part with common stock options. Pursuant to the merger and reorganization agreement between Abreos Biosciences, Inc. and Fimafeng, Inc., W.M.Y. surrendered all of his common stock options in 03/2021. **J.E. Maxson** discloses a collaboration with Ionis pharmaceuticals, research funding from Gilead Sciences, research funding from Kura Oncology and research funding from Blueprint Medicines. **J.W. Tyner** has received research support from Acerta, Agios, Aptose, Array, AstraZeneca, Constellation, Genentech, Gilead, Incyte, Janssen, Kronos, Meryx, Petra, Schrodinger, Seattle Genetics, Syros, Takeda, and Tolero. **B.J. Druker** potential competing interests-- SAB: Adela Bio, Aileron Therapeutics, Therapy Architects (ALLCRON), Cepheid, Celgene, DNA SEQ, Nemucore Medical Innovations, Novartis, RUNX1 Research Program, Vivid Biosciences (inactive); SAB & Stock: Aptose Biosciences, Blueprint Medicines, Enliven Therapeutics, Iterion Therapeutics, GRAIL, Recludix Pharma; Board of Directors & Stock: Amgen, Vincerx Pharma; Board of Directors: Burroughs Wellcome Fund, CureOne; Joint Steering Committee: Beat AML LLS; Advisory Committee: Multicancer Early Detection Consortium; Founder: VB Therapeutics; Sponsored Research Agreement: Enliven Therapeutics, Recludix Pharma; Clinical Trial Funding: Novartis, Astra-Zeneca; Royalties from Patent 6958335 (Novartis exclusive license) and OHSU and Dana-Farber Cancer Institute (one Merck exclusive license, one CytoImage, Inc. exclusive license, and one Sun Pharma Advanced Research Company non-exclusive license); US Patents 4326534, 6958335, 7416873, 7592142, 10473667, 10664967, 11049247. **T.P. Braun** has received research support from Astra- Zeneca, Blueprint Medicines as well as Gilead Sciences and is the institutional PI on the FRIDA trial sponsored by Oryzon Genomics. The authors certify that all compounds tested in this study were chosen without input from any of our industry partners. The other authors do not have competing interests, financial or otherwise.

## AUTHORS’ CONTRIBUTIONS

**W.M. Yashar**: conceptualization, software, formal analysis, validation, investigation, visualization, methodology, writing-original draft, writing-review and editing. **B.M. Smith**: conceptualization, software, formal analysis, validation, investigation, visualization, methodology, writing-review and editing. **D.J. Coleman**: conceptualization, formal analysis, validation, investigation, visualization, methodology. **J. VanCampen**: software. **G. Kong**: software. **J. Macaraeg**: validation, investigation, methodology. **J. Estabrook**: software. **E. Demir**: software. **N. Long**: data curation, writing-review and editing. **D. Bottomly**: data curation. **S.K. McWeeney**: data curation. **J.W. Tyner**: resources, data curation, supervision, funding acquisition, writing-review and editing. **B.J. Druker**: conceptualization, resources, supervision, funding acquisition, writing-review and editing. **J.E. Maxson**: conceptualization, resources, supervision, funding acquisition, writing-review and editing. **T.P. Braun:** conceptualization, resources, formal analysis, supervision, funding acquisition, validation, investigation, visualization, methodology, writing-original draft, project administration, writing-review and editing. The co-first authors may identify themselves as lead authors in their respective CVs.

## ACKNOWLEDGEMENTS

We would like to thank all of the BeatAML patients for their precious time and donation of samples supporting this research. We appreciate the following OHSU core facilities for their assistance: Advanced Light Microscopy, Flow Cytometry Shared Resource, Massive Parallel Sequencing Shared Resource, ExaCloud Cluster Computational Resource, and the Advanced Computing Center.

## SUPPLEMENTARY FIGURES

**Supplementary Fig. 1:**
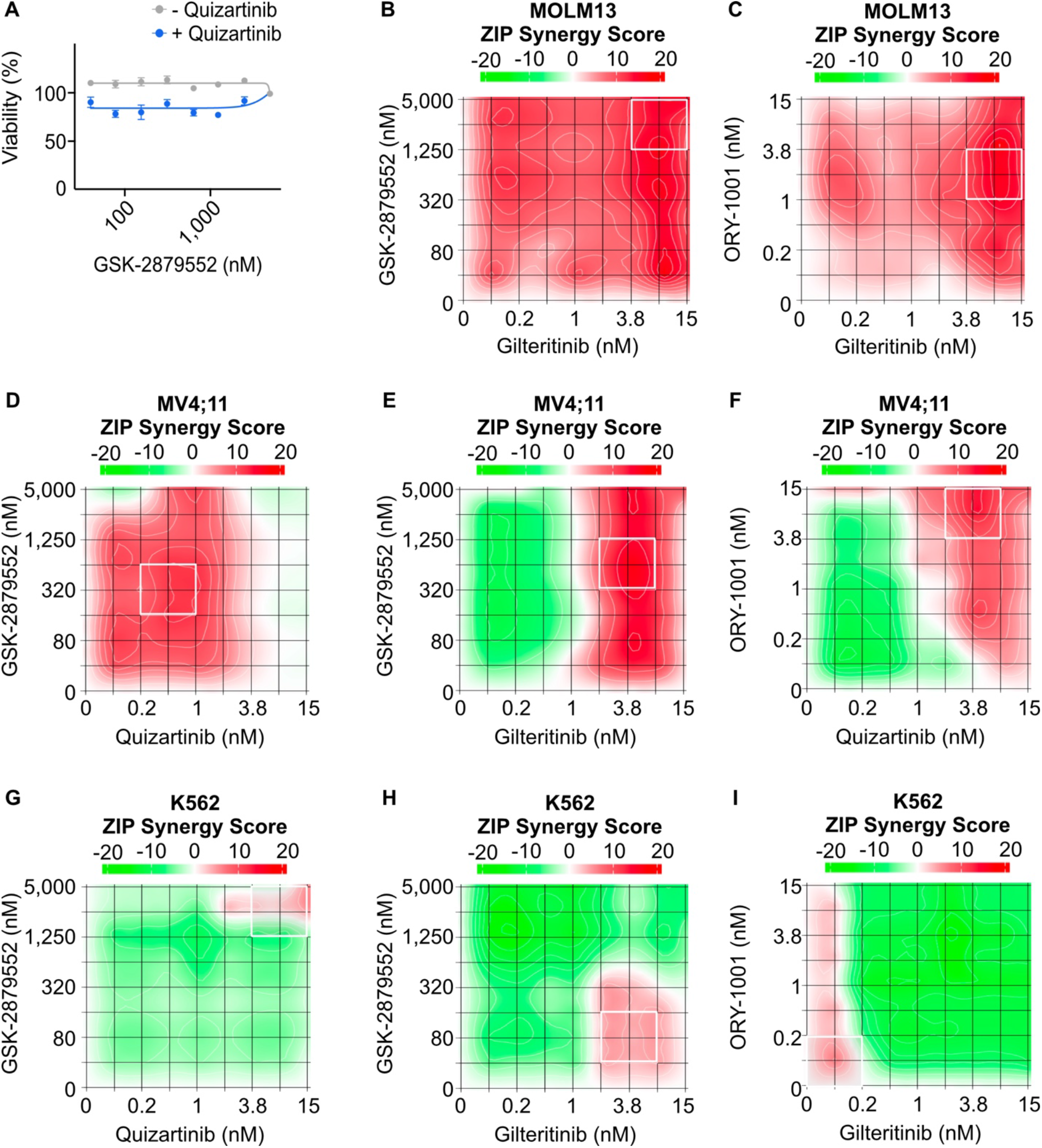
Drug synergy between FLT3 and LSD1 inhibitors in FLT3-ITD and FLT3 wild type cell lines. **A**, Dose response curves for GSK-2879552 with and without quizartinib (1 nM, which is the concentration corresponding to maximal synergy in Fig. 1). **B**, MOLM13 (FLT3-ITD-positive) cells were treated with an 8×8 matrix of gilteritinib and GSK-2879552 or **(C**) gilteritinib and ORY-1001 in triplicate. **D**, MV4;11 (FLT3-ITD-positive) cells were treated with an 8×8 matrix of quizartinib and GSK-2879552, **(E**) gilteritinib and GSK-2879552, or (**F)** quizartinib and ORY-1001 in triplicate. **G**, K562 (FLT3-wild-type) cells were treated with an 8×8 matrix of quizartinib and GSK-2879552, (**H**) gilteritinib and GSK-2879552, or (**I**) gilteritinib and ORY-1001 in triplicate. Cell viability was assessed after 3 days of culture using CellTiter Aqueous colorimetric assay. Synergy was assessed using the ZIP method.

**Supplementary Fig. 2:**
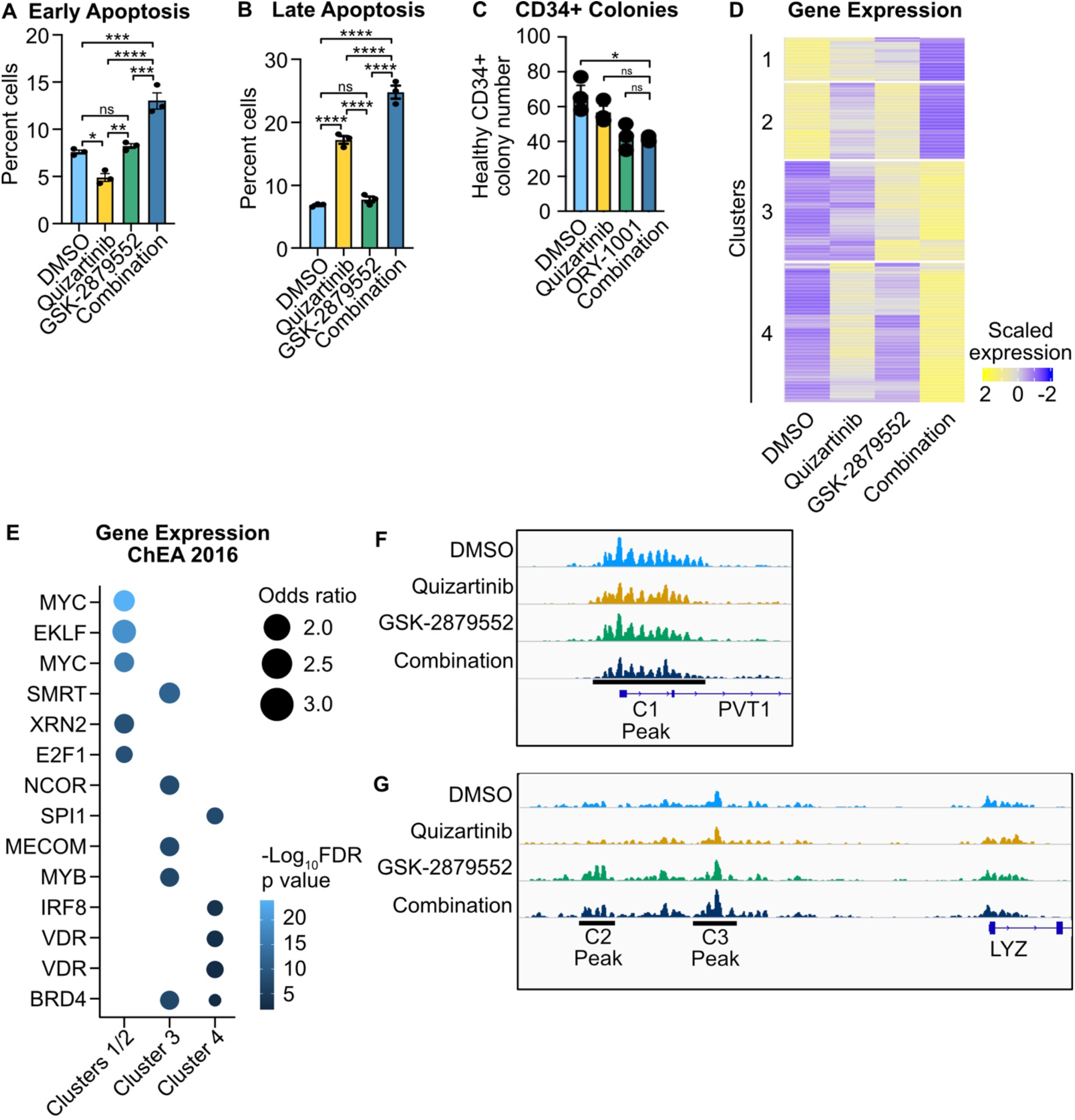
Efficacy of dual FLT3 and LSD1 inhibition in MOLM13 cells. **A, B**, MOLM 13 cells were treated quizartinib (1 nM), GSK-2879552 (100 nM), the combination, or an equal volume of DMSO vehicle for 48 hours prior to assessment of apoptosis by flow cytometry. Early apoptosis is Annexin V positive and propidium iodide negative. Late apoptosis is Annexin V positive and propidium iodide positive. Statistical significance was determined by two-way ANOVA with a Holm-Šidák post-test correction. **C**, CD34+ cells from healthy donor marrow were plated in complete human methocult media along with quizartinib (1 nM), GSK-2879552 (100 nM) the combination, or an equal volume of DMSO. Colony number was assessed after 14 days of growth. Statistical significance was determined by two-way ANOVA with a Holm-Šidák post-test correction. **D**, Unsupervised hierarchical clustering of differentially expressed genes and **E**, complete transcription factor target enrichment from RNA-seq on MOLM13 cells performed in Fig. 1B. **F, G**, Example tracks from H3K27ac CUT&Tag described in Fig. 1E. Examples of different clusters of differential peaks are shown. ns = not significant, * = p < 0.05, ** = p < 0.01, *** = p < 0.001, **** = p < 0.0001.

**Supplementary Fig. 3:**
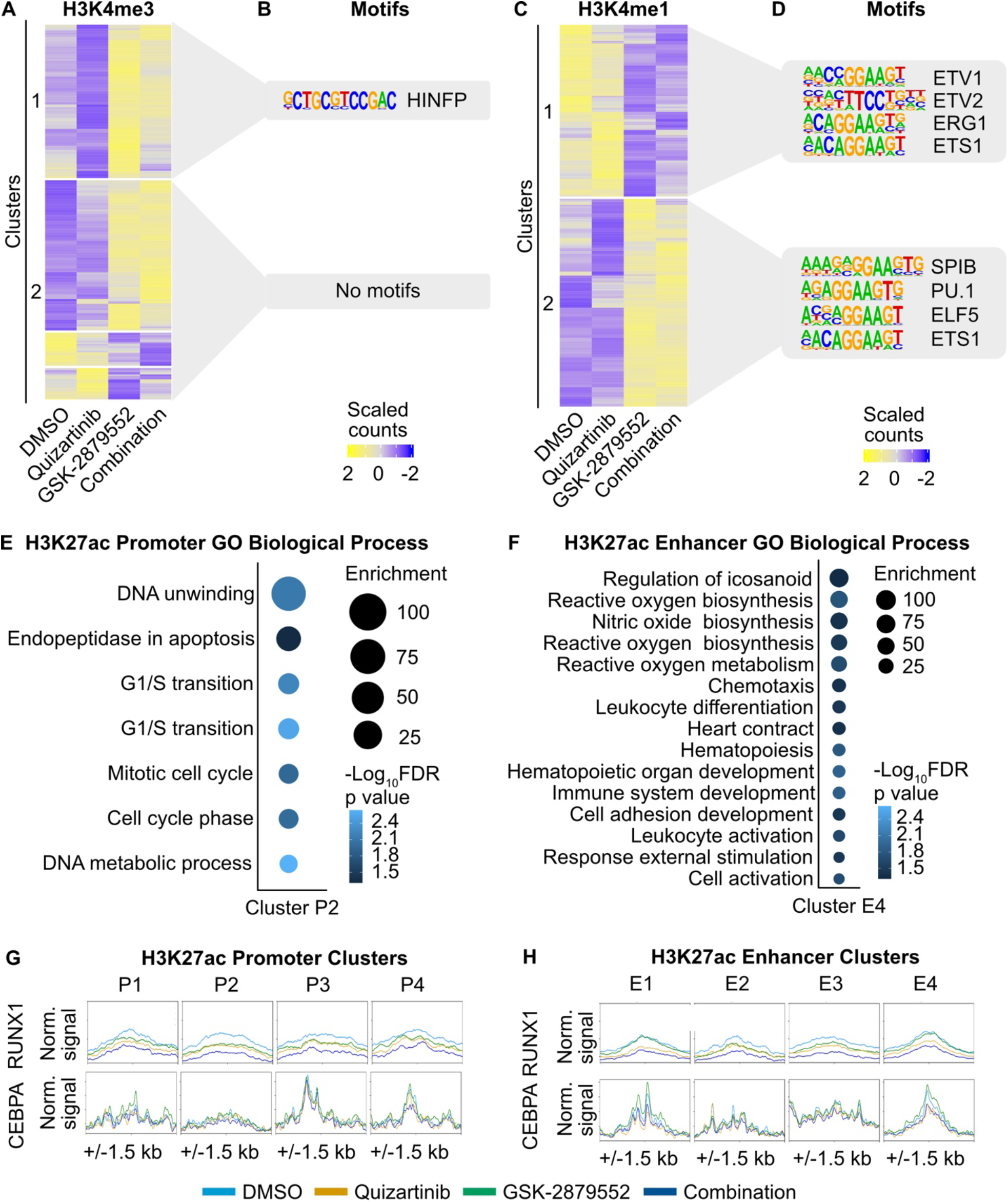
Epigenetic impact of dual FLT3 and LSD1 inhibition. **A**, Unsupervised hierarchical clustering of regions of differential H3K4me3 at promoters assessed by CUT&Tag from MOLM13 cells treated with quizartinib (1 nM), GSK-2879552 (100 nM), the combination, or an equal volume of DMSO for 6 hours. **B**, Motif enrichment for regions of differential H3K4me3 read density at promoters. Top 4 enriched transcription factor motifs are shown. **C**, Unsupervised hierarchical clustering of regions of differential H3K4me1 at enhancers assessed by CUT&Tag from MOLM13 cells treated with quizartinib (1 nM), GSK-2879552 (100 nM), the combination, or an equal volume of DMSO for 6 hours. **D**, Motif enrichment for regions of differential H3K4me1 read density at enhancers. Top 4 enriched transcription factor motifs are shown. **E, F**, GO analysis of differential H3K27ac read pileup at (**E**) promoters and (**F**) enhancers from analysis in Fig. 2. Dot size represents the binomial fold enrichment and color represents the −log_10_(FDR p-value) of the GO Biological Process term. **G**, MOLM13 cells were treated with quizartinib (1 nM), GSK-2879552 (100 nM), the combination, or an equal volume of DMSO for 6 hours. RUNX1 binding was assessed by ChIP-seq and CEBPA binding was assessed by CUT&Tag. Transcription factor binding profiles at promoters with differential H3K27ac identified in Fig. 2. **H**, Transcription factor profiles at enhancers with differential H3K27ac identified in Fig. 2.

**Supplementary Fig. 4:**
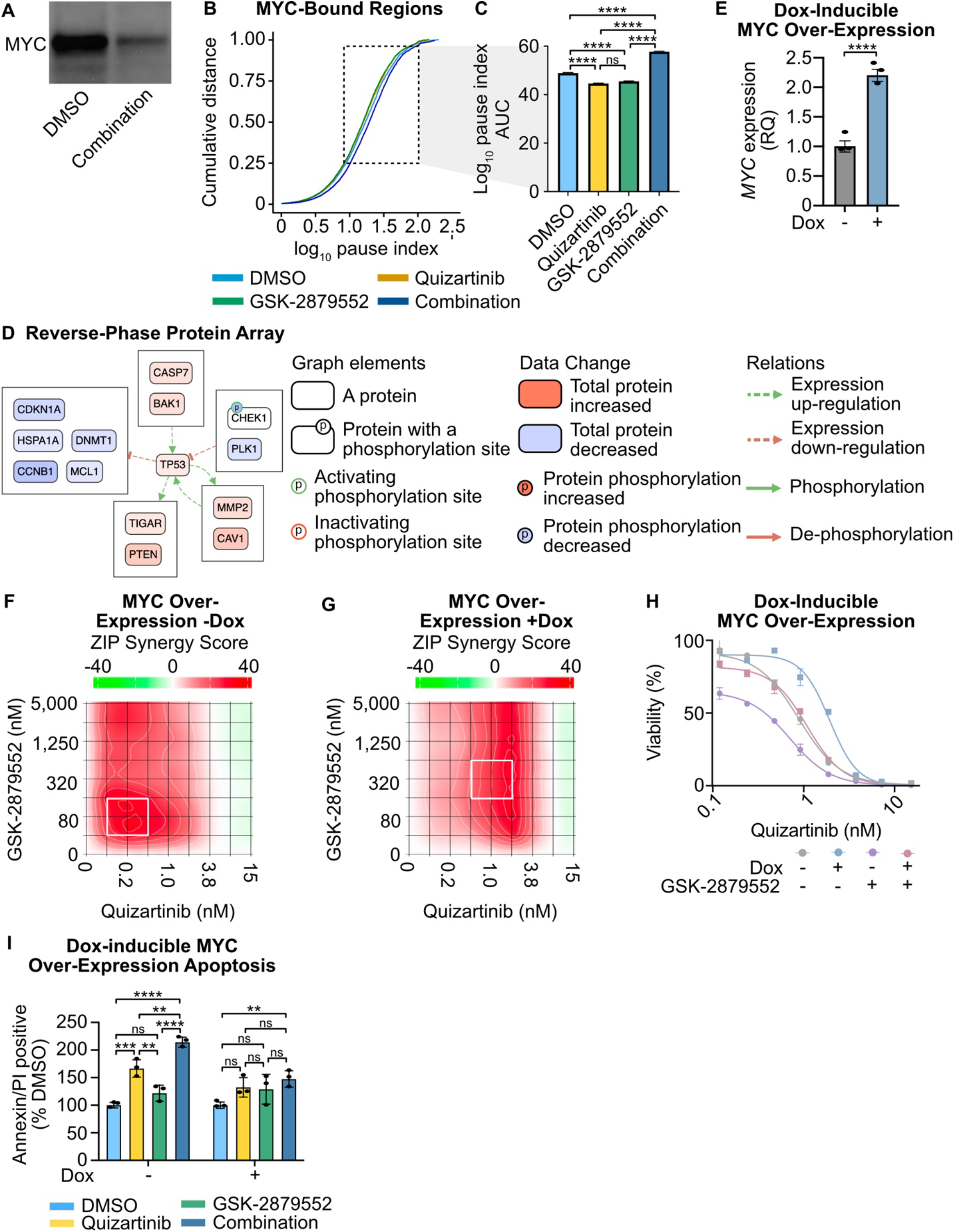
Dual FLT3 and LSD1 inhibition suppresses *MYC* expression and activity. **A**, Western blot for MYC in MOLM13 cells treated with dual quizartinib (1 nM) and GSK-2879552 (100 nM) or an equivalent volume of DMSO for 24 hours. **B**, MOLM13 cells were treated with quizartinib (1 nM), GSK-2879552 (100 nM), the combination, or an equal volume of DMSO vehicle for 6 hours prior to CUT&Tag for RBP1. Cumulative distribution of RNA PolII pause-indices at RBP1 and MYC co-bound genes. **C**, AUC of RNA PolII pause-indices within the 25-95% cumulative distribution. Statistical significance was determined by Kolmogorov-Smirnov test with an FDR post-test correction. **D**, Differential phosphoprotein network enrichment in MOLM13 cells treated for 24 hours with 1 nM quizartinib and 100 nM GS-K2979552 or an equivalent volume of DMSO. **E**, MOLM13 cells were transduced with lentiviral particles harboring a doxycycline-inducible *MYC* expression vector. *MYC* expression was assessed 48 hours after the addition of doxycycline (1 µg/mL) and normalized to *GAPDH* as an endogenous control. Statistical significance was determined by Student’s t-test. **F, G**, Cells were treated doxycycline (1 µg/mL) or an equivalent volume of DMSO for 72 hours then plated in triplicate in an 8×8 matrix of concentrations of quizartinib and GSK-2879552 for 3 days. Cell viability was measured using CellTiter Aqueous colorimetric assay. Synergy was assessed using the ZIP method. **H**, Quizartinib response curves in MOLM13 cells harboring a doxycycline-inducible *MYC* expression construct. Cells were also treated with and without doxycycline (1 µg/mL) and/or GSK-2979552 (311 nM, which is the concentration corresponding to maximal synergy in the *MYC* over-expressed MOLM13 cells in g). **I**, Cells were treated for 72 hours with doxycycline (1 µg/mL) then treated with quizartinib (1 nM), GSK-2879552 (100 nM), the combination, or an equal volume of DMSO for 48 hours. Apoptosis was assessed using flow cytometry for Annexin V and PI. Statistical significance was determined by two-way ANOVA with a Holm-Śidák post-test correction. ns = not significant, * = p < 0.05, ** = p < 0.01, *** = p < 0.001, **** = p < 0.0001.

**Supplementary Fig. 5:**
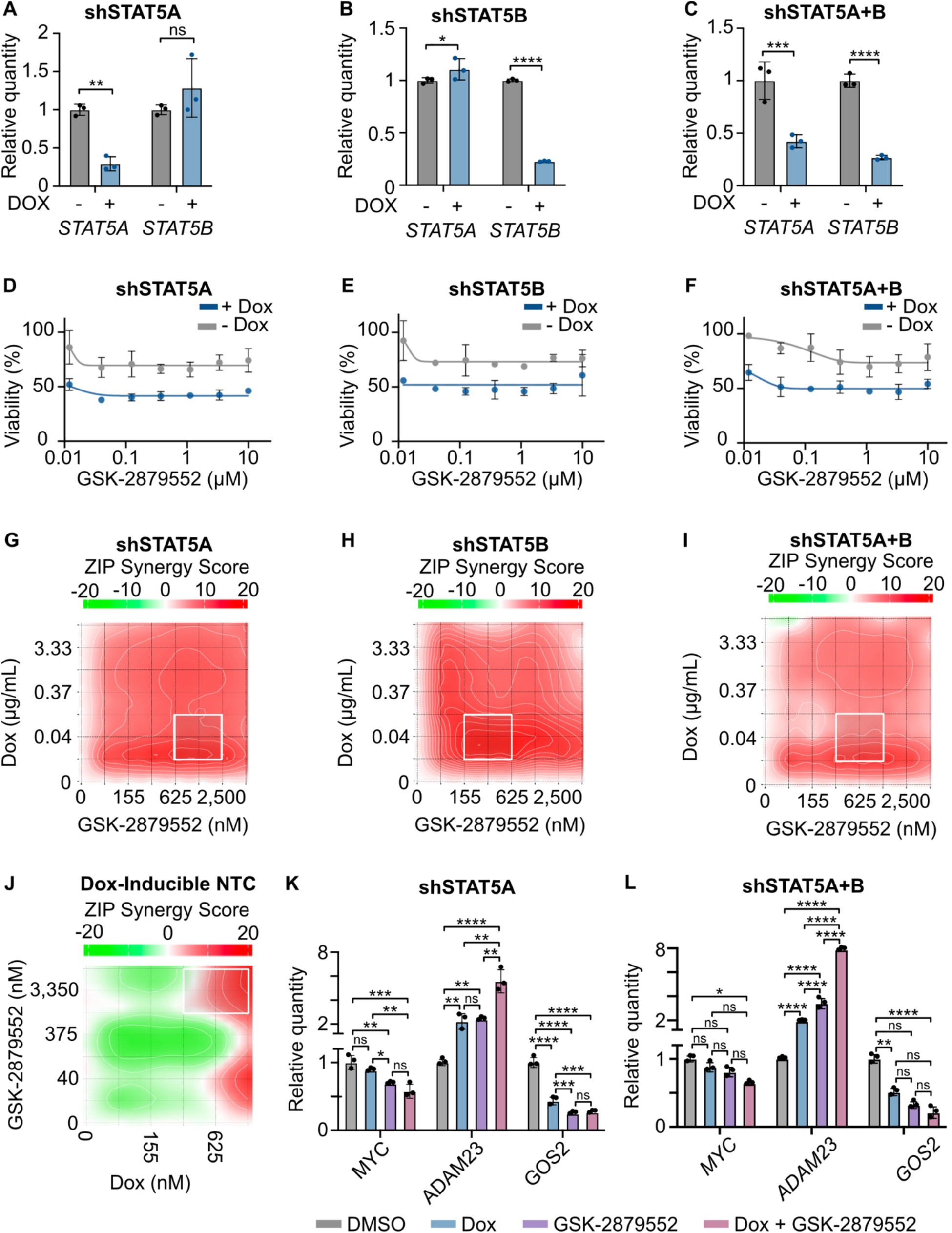
STAT5 and MYC play a key role in the response to FLT3 and LSD1 inhibition. **A-C**, MOLM13 cells were transduced with lentiviral particles harboring a doxycycline-inducible *STAT5* knockdown. Cells were induced with doxycycline (1 µg/mL) for 48 hours. Validation of knockdown of *STAT5A* and *STA5B* by qPCR relative to *GUSB* endogenous control. Significance was determined by Student’s t-test. **D-F**, Dose response curves for GSK-2879552 with and without doxycycline (1 µg/mL). **G, H**, 8×8 drug synergy matrices between GSK-2879552 and (**G**) *STAT5A*, (**H**) *STAT5B*, or (**I**) *STAT5A* and *STAT5B* knockdown (induced by doxycycline) were performed in triplicate in MOLM13 cells with viability assessed by CellTiter Aqueous colorimetric assay after 3 days of drug exposure. ZIP synergy scores were calculated on the average values for each drug dose. **J**, MOLM13 cells harboring a doxycycline-inducible non-targeting control (iNTC) shRNA construct were treated doxycycline (1 µg/mL) or an equivalent volume of DMSO for 72 hours then plated in triplicate in an 8×8 matrix of concentrations of GSK-2879552 and doxycycline for 3 days. Cell viability was measured using CellTiter Aqueous colorimetric assay. Synergy was assessed using the ZIP method. **K, L**, qPCR assessment of gene expression in MOLM13 cells expressing a doxycycline-inducible (**K**) *STAT5A* or (**L**) *STAT5A* and *STAT5B* shRNA. Cells were treated with doxycycline (1 µg/mL) or an equivalent volume of DMSO for 72 hours prior to the addition of GSK-2879552 (100 nM) for 24 hours. Expression was normalized to *GusB* as an endogenous control. Statistical significance was determined by two-way ANOVA with a Holm-Šidák post-test correction. ns = not significant, * = p < 0.05, ** = p < 0.01, *** = p < 0.001, **** = p < 0.0001.

**Supplementary Fig. 6:**
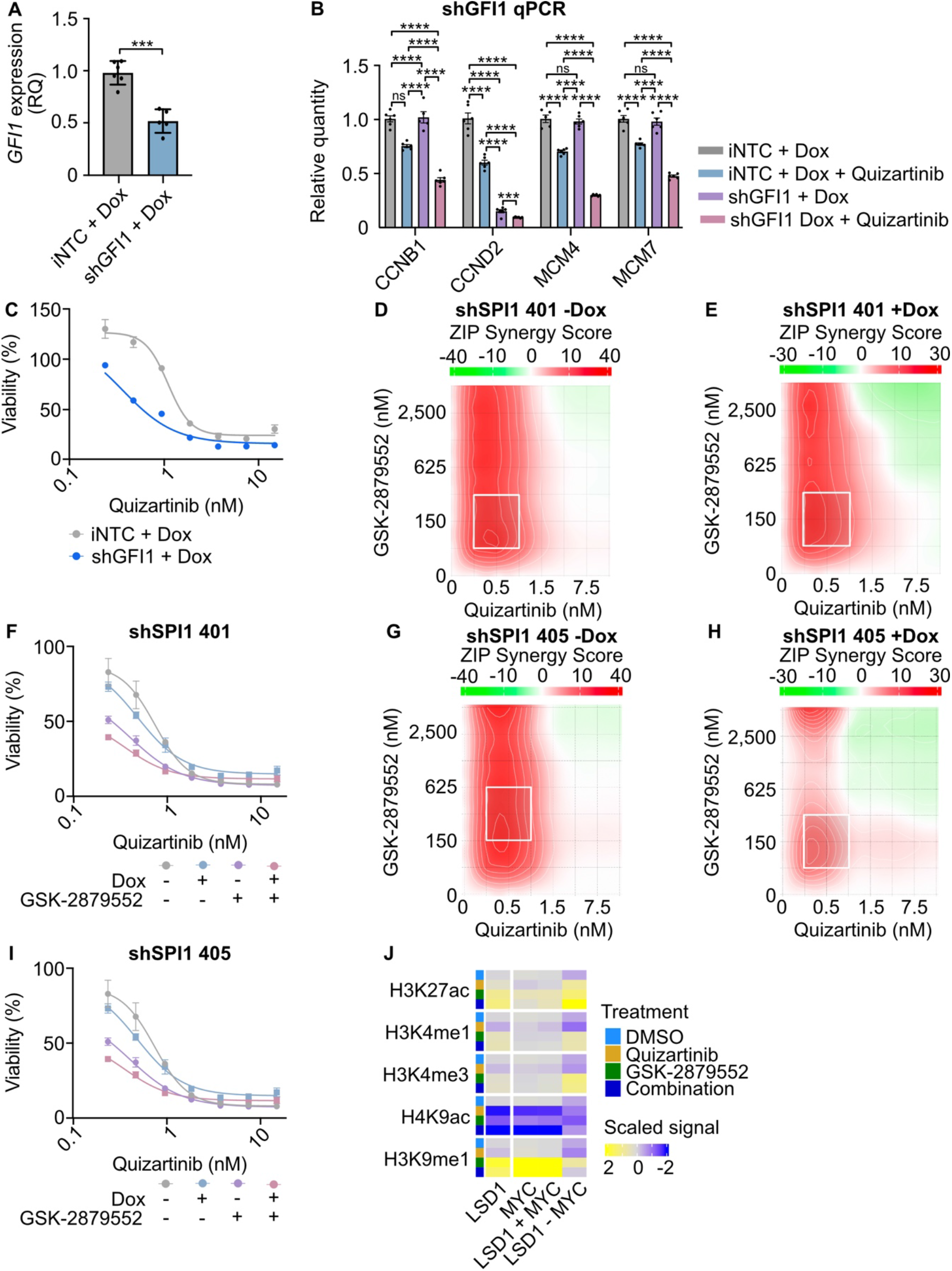
Knockdown of GFI1 weakens the effects of dual FLT3 and LSD1 inhibition. **A**, Validation of *GFI1* knockdown by qPCR relative to *GUSB* endogenous control. Statistical significance was determined by Student’s t-test. **B**, qPCR assessment of gene expression in MOLM13 cells expressing a doxycycline-inducible *GFI1* shRNA. Cells were treated with doxycycline (1 µg/mL) for 72 hours prior to the addition of quizartinib (1 nM) for 24 hours. Substantial knockdown was observed in the absence of doxycycline treatment, so only doxycycline-treated samples were compared. Expression was normalized to *GusB* as an endogenous control. Statistical significance was determined by two-way ANOVA with a Holm-Śidák post-test correction. **C**, Dose response curves for MOLM13 cells harboring a doxycycline-inducible iNTC or *GFI1* shRNA with doxycycline (1 µg/mL). **D, E**, 8×8 drug synergy matrices between GSK-2879552 and quizartinib with and without *SPI1* shRNA construct 401 (induced by doxycycline) were performed in triplicate in MOLM13 cells with viability assessed by CellTiter Aqueous colorimetric assay after 3 days of drug exposure. **F**, Dose response curves in MOLM13 cells harboring a doxycycline-inducible *SPI1* shRNA construct 401 with quizartinib. Cells were also treated with and without doxycycline (1 µg/mL) and/or GSK-2979552 (311 nM, which is the concentration corresponding to maximal synergy in the *SPI1* knockdown MOLM13 cells in e). **G, H**, Same as (**D**) and (**E**) except with the *SPI1* shRNA construct 405. **I**, Same as (**F**) except with the *SPI1* shRNA construct 405. **J**, MOLM13 cells were treated with quizartinib (1 nM), GSK-2879552 (100 nM), the combination, or an equal volume of DMSO for 6 hours before histone modification assessment. H3K9ac and H3K9me1 binding were assessed using CUT&Tag. Heatmap depiction of normalized read pileup of H3K9ac and H3K9me1 as well as H3K27ac, H3K4me1, and H3K4me3 at LSD1 and MYC binding sites (data taken from Fig. 2 excluding H3K9 marks). ns = not significant, * = p < 0.05, ** = p < 0.01, *** = p < 0.001, **** = p < 0.0001.

**Supplementary Fig. 7:**
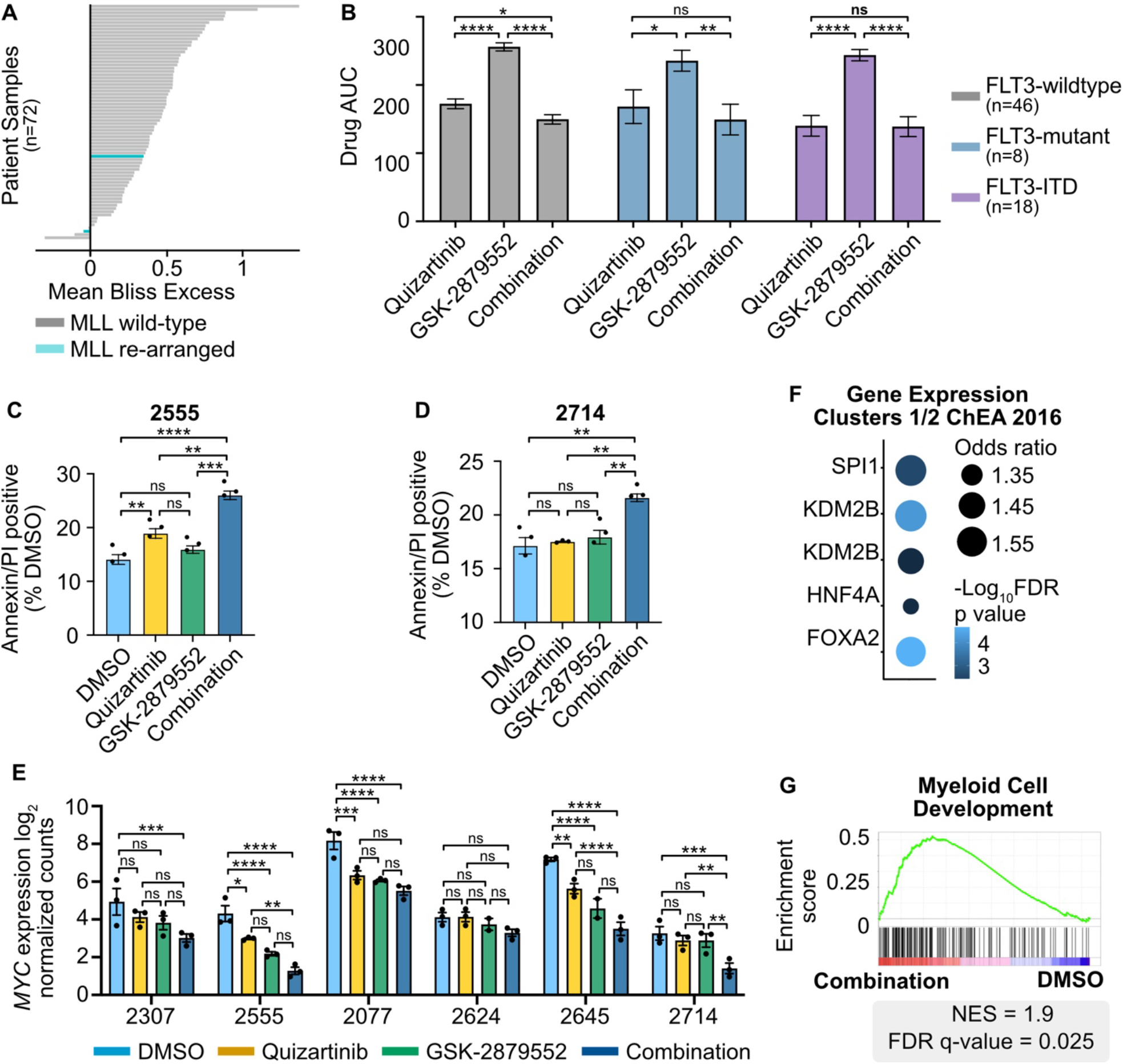
Efficacy of dual FLT3 and LSD1 inhibition in primary AML samples. **A**, Primary AML blasts from 72 total samples (18 FLT3-ITD-positive) were cultured for 3 days along a 7-point curve with either quizartinib, GSK-2879552, or equimolar amounts of the drug combination. Cell viability was assessed by CellTiter Aqueous colorimetric assay. The excess over bliss score was calculated using cell viability at corresponding drug concentrations. Each bar represents the mean excess over bliss across all concentrations. Bar color indicates MLL re-arrangement status. **B**, Mean drug AUC in patient samples grouped by FLT3 mutation status. Statistical significance was determined by two-way ANOVA with a Holm-Śidák post-test correction. **C, D**, Primary AML blasts were treated with quizartinib (500 nM), GSK-2879552 (500 nM), the combination, or an equivalent volume of DMSO for 24 hours. Apoptosis was assessed by flow cytometry for Annexin V and PI after 48 hours of drug exposure. Statistical significance was determined by two-way ANOVA with a Holm-Śidák post-test correction. **E**, RNA-seq was performed on six independent, FLT3-ITD-positive patient samples treated in triplicate with 500 nM quizartinib, 500 nM GSK-2879552, both drugs in combination, or an equivalent volume of DMSO for 24 hours. *MYC* gene expression in each individual sample is shown. Statistical significance was determined by two-way ANOVA with a Holm-Šidák post-test correction. **F**, Transcription factor target enrichment from clusters in Fig. 6G. **G**, Select GSEA plot from clusters in Fig. 6G. ns = not significant, * = p < 0.05, ** = p < 0.01, *** = p < 0.001, **** = p < 0.0001.

**Supplementary Fig. 8:**
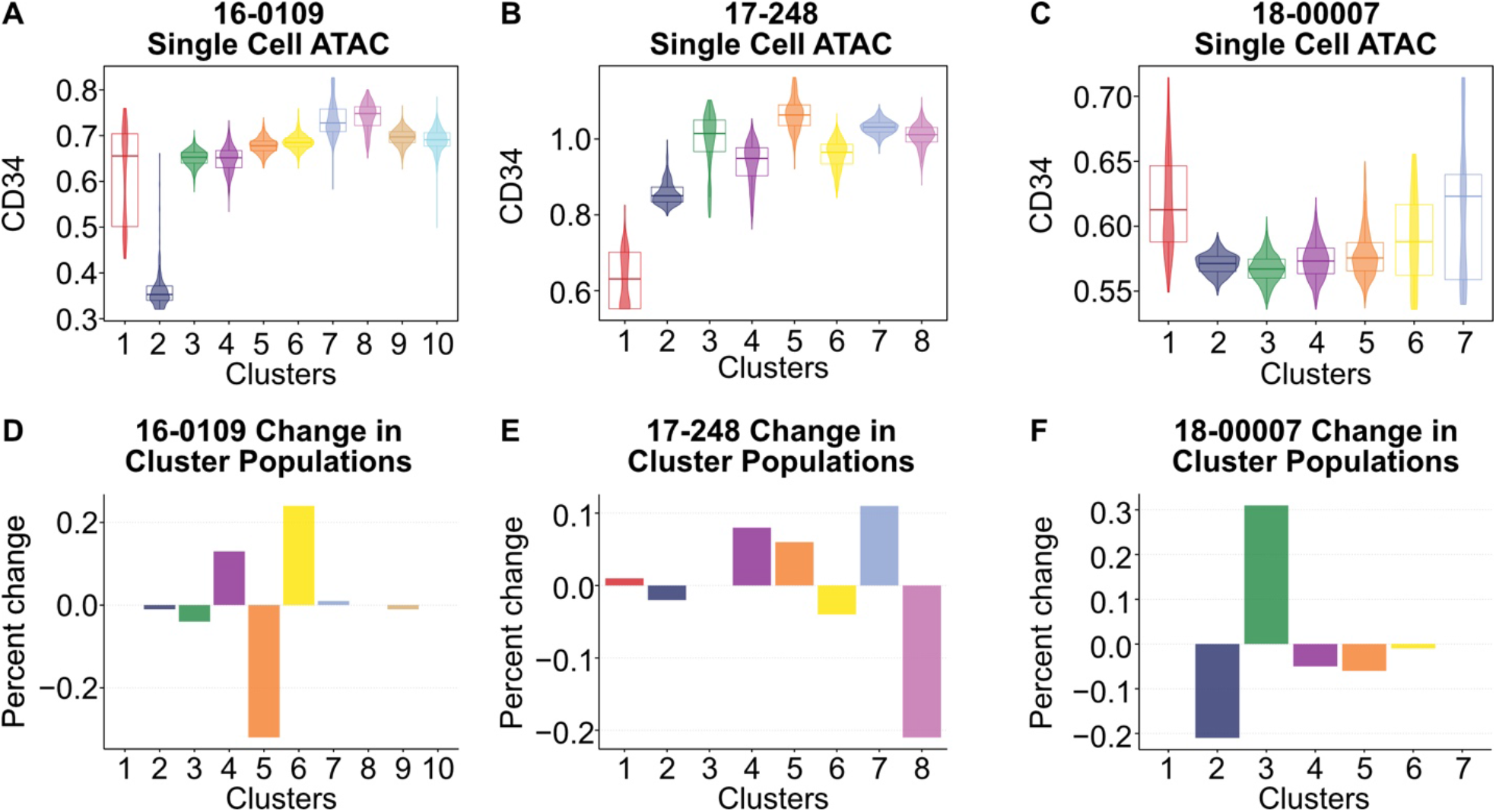
Marker for immature populations in primary patient sample single cell ATAC-seq. **A-C**, Primary AML blasts from three FLT3-ITD-positive samples MOLM13 cells were treated with quizartinib (500 nM) and GSK-2879552 (500 nM) or an equal volume of DMSO vehicle for 24 hours prior to single cell ATAC sequencing. Clusters contain cell from both conditions. The CD34 gene scores were calculated from chromatin accessibility and are an estimate of gene transcription. **D-F**, Percent change in cluster populations between DMSO and combination treatments.

## Notes

### Summary of Updates

Figures and paper structure

